# Role of the Topoisomerase IIα Chromatin Tether domain in Nucleosome Binding & Chromosome Segregation

**DOI:** 10.1101/2021.10.03.462934

**Authors:** Sanjana Sundararajan, Hyewon Park, Shinji Kawano, Marnie Johansson, Tomoko Saito-Fujita, Noriko Saitoh, Alexei Arnaoutov, Mary Dasso, Duncan J. Clarke, Yoshiaki Azuma

## Abstract

Due to the intrinsic nature of DNA replication, replicated genomes retain catenated genomic loci that must be resolved to ensure faithful segregation of sister chromatids in mitosis. Type II DNA Topoisomerase (TopoII) decatenates the catenated genomic DNA through its unique Strand Passage Reaction (SPR). Loss of SPR activity results in anaphase chromosome bridges and formation of <u>P</u>olo-like Kinase <u>I</u>nteracting <u>C</u>heckpoint <u>H</u>elicase (PICH)-coated ultra-fine DNA bridges (UFBs) whose timely resolution is required to prevent micronuclei formation. Vertebrates have two TopoII isoforms– TopoIIα and TopoIIβ, that share a conserved catalytic core. However, the essential mitotic function of TopoIIα cannot be compensated by TopoIIβ, due to differences in their catalytically inert C-terminal domains (CTDs). Using genome-edited human cells, we show that specific binding of TopoIIα to methylated histone, tri-methylated H3K27 (H3K27me3), via its Chromatin Tether (ChT) domain within the CTD contributes critically to avoid anaphase UFB formation. Reducing H3K27 methylation prior to mitosis increases UFBs, revealing a requirement for proper establishment of H3K27me3 after DNA replication to facilitate TopoIIα-ChT dependent UFB prevention. We propose that interaction of the TopoIIα-ChT with H3K27me3 is a key factor that ensures the complete resolution of catenated loci to permit faithful chromosome segregation in human cells.

**Summary Statement:** Genomic catenations originating from the DNA replication process must be resolved by DNA Topoisomerase II (TopoII) to permit sister chromatid disjunction. The results show that specific recognition of methylated histone containing chromatin by TopoII is critical for complete resolution of the genome.

## Introduction

During mitosis, the products of DNA replication must be meticulously partitioned between the two daughter cells. Towards this pursuit, the sister chromatids that exist in a highly catenated state, due to the intrinsic nature of DNA, can only be resolved by Type II DNA Topoisomerases that can decatenate DNA using their Strand Passage Reaction (SPR) (Nitiss, 2009). Vertebrates express two isoforms of Topoisomerase II (TopoIIα and TopoIIβ), encoded by separate genes and distinguished by their molecular weight. Owing to their conserved ATPase domains and catalytic cores, they show similar SPR activity in *in vitro* decatenation assays (Gilroy and Austin, 2011). However, their C-terminal Domains (CTDs), that show much lower homology, distinguish the isoforms and account for the diversity in their cellular functions; TopoIIα is essential for chromosome condensation and segregation, whereas TopoIIβ is dispensable for mitosis, but has crucial roles in regulating transcription, particularly in non-dividing cells (Grue et al., 1998; Linka et al., 2007; Sakaguchi and Kikuchi, 2004).

Although it is well established that TopoIIα is a major component of mitotic chromatin and its activity is necessary for proper mitotic progression (Earnshaw et al., 1985; Grue et al., 1998; Nielsen et al., 2020), we do not know the exact mechanisms that account for this and distinguish TopoIIα from TopoIIβ. In addition, little is known about how TopoIIα is specifically recruited to catenated loci that must be resolved before anaphase. Indeed, for a long time it was difficult to clearly visualize the remaining catenated loci in mitosis that TopoII must target. The discovery of PLK-1 Interacting Checkpoint Helicase (PICH) revolutionized the study of the catenated genome (Baumann et al., 2007). PICH localizes to catenated loci that persist in anaphase, after sister chromatids begin to segregate. Most of these catenations are between DNA molecules of sister centromeres (CENs), are referred to as anaphase Ultra-Fine DNA Bridges (UFBs) and must be efficiently targeted for resolution by late anaphase (Biebricher et al., 2013). Even small numbers of residual UFBs gives rise to micronuclei that accumulate damaged DNA following cytokinesis (Hengeveld et al., 2015). Hence, with persistence of unresolved catenanes, there is an accumulation of genomic lesions that can contribute to tumorigenesis (Krupina et al., 2021).

Previous studies indicated that siRNA knockdown of TopoIIα but not TopoIIβ results in increased UFBs (Antoniou-Kourounioti et al., 2019; Spence et al., 2007). This reflects the function of TopoIIα in resolving these catenated loci prior to anaphase. However, we do not know what exact mechanisms are involved in targeting TopoIIα to the catenations, i.e., to the anaphase UFB precursors. Bulk decatenation of the genome may be efficiently achieved stochastically, but when a small number of CEN catenations remains, a highly targeted mechanism presumably exists to ensure their decatenation. Molecular insight into this mechanism has emerged from study of the TopoIIα Chromatin Tether (ChT) domain, which comprises the terminal 31aa of TopoIIα and, *in vitro,* binds to methylated Histone H3 (Lane et al., 2013). Binding assays with modified histone H3 N-terminal peptides demonstrated that the ChT domain binds to peptides methylated at lysine 27 and arginine 26 whereas phosphorylation at serine 28 repulses this binding. Cells depleted of both TopoII isoforms and expressing a mutant TopoIIα lacking the ChT (TopoIIα–ΔChT) exhibited mitotic defects; 1) a defect in sister chromatid resolution/individualization, and 2) reduced condensation (Lane et al., 2013). TopoIIα–ΔChT has a higher dissociation rate from mitotic chromosomes suggesting that the ChT domain is required for the optimal residence time of TopoIIα on mitotic chromosomes. These results hint that the *in vitro* interaction of the ChT with histones is biologically important *in vivo*. However, whether the ChT binds to nucleosomes with methylated histones *in vivo* has not been determined, and it remains unknown if such epigenetic cues control TopoIIα function on mitotic chromosomes to facilitate efficient genome resolution.

Here, we used the Auxin Inducible Degron (AID) system coupled with Doxycycline inducible (Tet-ON) gene replacement in DLD-1 cells to reveal the critical function of the TopoIIα ChT domain (αChT) in prevention of UFBs. We show that the αChT is required for binding to H3K27me3 mononucleosomes *in vitro* and for complete prevention of UFBs *in vivo;* these functions distinguish TopoIIα from TopoIIβ. Inhibition of H3K27me3 by the potent methyltransferase inhibitor, GSK-343, increased the frequency of UFBs, consistent with loss of αChT function. The composition of aromatic amino acids within the αChT is also critical for both H3K27me3 binding and αChT-mediated prevention of UFBs, consistent with the possibility that TopoIIα uses an aromatic cage structure for chromatin binding, similar to the mechanism that has been proposed for other chromatin proteins that interact with methylated lysines (Jacobs and Khorasanizadeh, 2002; Min et al., 2003; Nielsen et al., 2002). Together, we demonstrate that H3K27me3 is a novel epigenetic cue employed by TopoIIα for the recognition and resolution of catenated loci to permit faithful chromosome segregation.

## Results

### TopoIIα, not TopoIIβ, is essential for the prevention of increased PICH positive UFBs

One of the defects previously observed in TopoIIα-depleted cells was increased UFBs in anaphase due to lack of resolution of catenated centromeric loci (Spence et al., 2007). Using our established AID-mediated TopoIIα-depletion cell lines (Hassebroek et al., 2020), we were able to quantify UFBs after TopoIIα-depletion within a single cell division cycle. In addition, we created AID-TopoIIβ cell lines for comparison (Supplemental fig. 1). Similar to AID-fused TopoIIα, AID-fused TopoIIβ is efficiently degraded upon Auxin (Aux) addition (Supplemental fig. 1). To analyze UFBs in anaphase, AID-TopoII cells were synchronized by a single thymidine block, Aux was added upon release, and mitotic cells were collected by shake-off after reaching mitosis (Fig. 1 A, see Materials and Methods). Degradation of the proteins in each case was confirmed by western blot using antibodies against TopoIIα/TopoIIβ and FLAG-tag, located between the AID and TopoII (Fig. 1 B). FLAG-tag signals showed that the amount of endogenous TopoIIα is much higher than that of endogenous TopoIIβ. Consistent with previous results (Hassebroek et al., 2020), chromosome fractions from TopoIIα-depleted cells showed increased PICH signals. In contrast, TopoIIβ-depletion did not increase PICH associated with mitotic chromosomes. Notably, the amount of chromosomal TopoIIβ was increased in TopoIIα-depleted chromosomes, suggesting that both isoforms might compete for chromosomal binding. To quantify UFBs in anaphase, cells were treated as in Fig. 1 A and then fixed and stained with antibodies against PICH, FLAG (for endogenous TopoII) and CENP-C (as a CEN marker) (Fig. 1 C and D). The AID-mediated TopoIIα-depleted cells had an increase in UFBs/cell as compared to cells with depleted TopoIIβ (Fig. 1 E). Our results not only recapitulate those from the previous study but also indicate that loss of TopoIIα in less than a single cell cycle, between S-phase and mitosis, results in a large increase in UFBs in anaphase.

**Figure 1.**
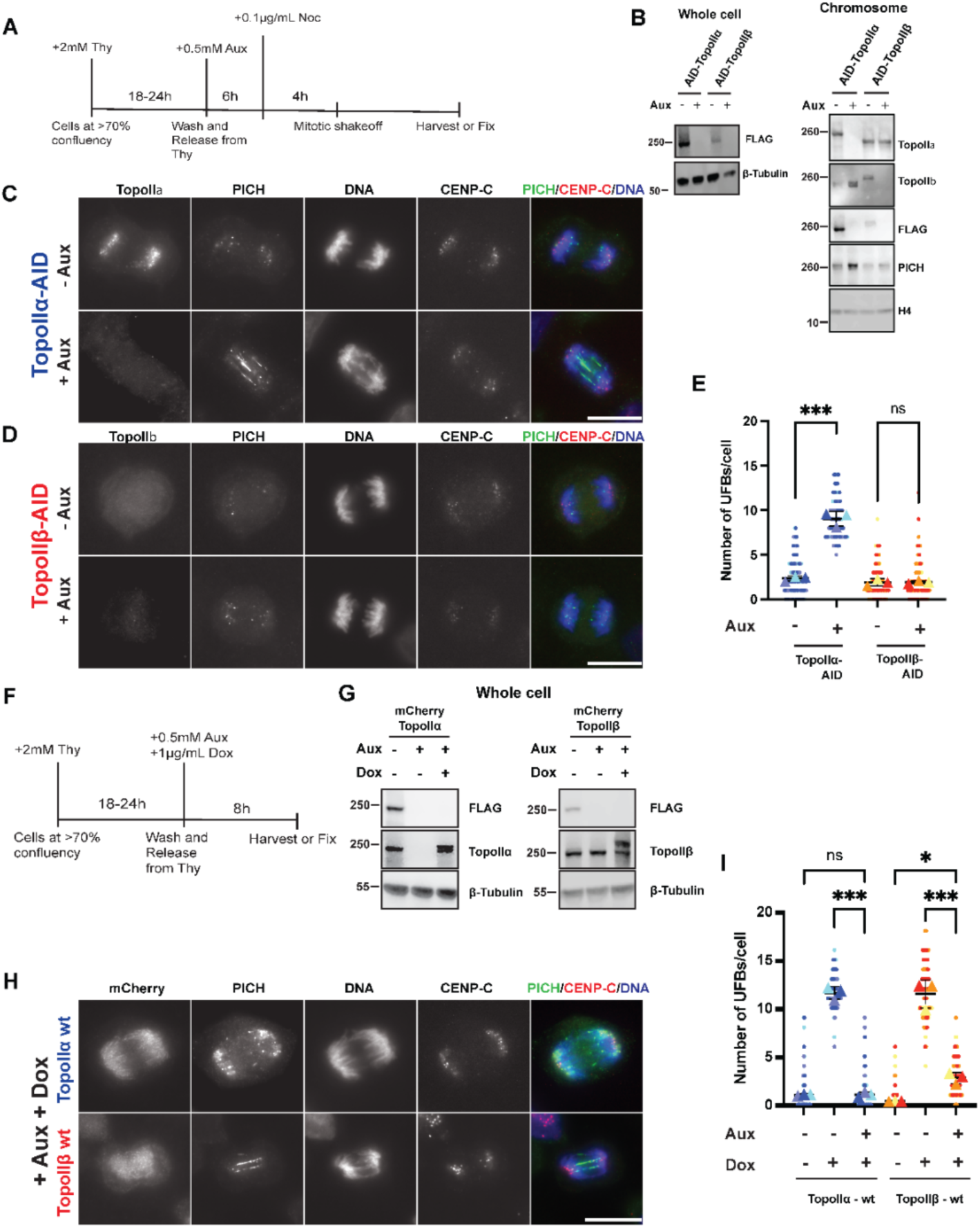
TopoIIα is required for preventing UFBs: **A)** Schematic of synchronized cell culture conditions for analyzing effect of TopoIIα and TopoIIβ degradation by AID system (for data in Fig 1B, 1C, 1D and 1E only). **B)** Western blot showing Aux induced depletion of TopoIIα and TopoIIβ (both tagged with FLAG) in engineered DLD-1 cells in the whole cell lysate (left) and on the chromosome (right) probed with antibodies against indicated targets. **C)** Increased number of UFBs observed in anaphase cells as indicated by staining with anti-PICH antibody, when TopoIIα is eliminated (bottom) as compared to presence of TopoIIα (top). CENs are marked with CENP-C. Bars, 10 μm. **D)** UFB analysis performed as in C with AID-TopoIIβ cells. TopoIIβ elimination does not result in an increase in UFBs (bottom) as compared to presence of TopoIIβ. Bars, 10 μm. **E)** Superplots showing quantification of the UFBs in terms of number of UFBs/cell (from >60 cells counted over three independent experiments) for TopoIIα (blue bullets) and TopoIIβ (red bullets). P-value indicates one-way ANOVA analysis followed by Tukey multicomparison correction for the means. Horizontal bars indicate mean and error bars indicate SD calculated for the means across the three independent experiments. ns: not statistically significant, ***: p<0.001. **F)** Schematic for synchronization method of cells with thymidine for all UFB assays. **G)** Western blot showing replacement of mCherry-TopoIIα (left) and mCherry-TopoIIβ (right) upon Dox addition in endogenous TopoIIα depleted cells. **H)** Representative Images acquired for UFB assay with either mCherry-TopoIIα (top) or mCherry-TopoIIβ (bottom) replacement. Bars, 10 μm. **I)** Superplots showing quantification of the number of UFBs/cell (from >60 cells counted over three independent experiments) for mCherry-TopoIIα (blue bullets) and mCherry-TopoIIβ (red bullets) replaced cells. P-value indicates one-way ANOVA analysis followed by Tukey multicomparison correction for the means. Horizontal bars indicate mean and error bars indicate SD calculated for the means across the three independent experiments. ns: not statistically significant, *: p<0.033, ***: p<0.001.

Although the results demonstrate a specific role of TopoIIα in resolving the catenated genome to prevent increased UFBs, the expression level of endogenous TopoIIβ is clearly lower than TopoIIα, and the chromosomal TopoIIβ was increased in TopoIIα-depleted cells (Fig. 1 B). This suggests that TopoIIβ might be able to compensate for the loss of TopoIIα if there is enough TopoIIβ available. Therefore, to further define what specificity there is between the isoforms for the prevention of UFBs, we designed rescue experiments using a Tet-inducible (Tet-ON) expression system. Tet-ON cassettes (Natsume et al., 2016) with mCherry-TopoIIα wild type (wt) or mCherry-TopoIIβ wt were integrated into the hH11 safe harbor loci as previously reported (Hassebroek et al., 2020) (Supplemental fig. 2). For these rescue experiments, cells were synchronized as described above to deplete TopoIIα, but with concurrent addition of Doxycycline (Dox) to induce the exogenous alleles of TopoII (Fig. 1 F). Western blot of whole cell lysates confirmed endogenous TopoIIα depletion and expression of the transgenes (Fig. 1 G). Immunofluorescence analysis to quantify UFBs in these cells showed that replacement with mCherry-TopoIIα efficiently restored the low abundance of UFBs to levels similar to the control. However, mCherry-TopoIIβ could not completely rescue the high frequency of UFBs seen in the depleted cells, even though mCherry-TopoIIβ was over-expressed; UFBs were significantly reduced compared to the TopoIIα-depleted cells but remained significantly more abundant compared to the control (Fig. 1 H and I). Incomplete restoration of normal UFB number in cells by over-expressed mCherry-TopoIIβ suggests that a subset of catenated loci require TopoIIα for their complete resolution.

### TopoIIα’s ability to prevent UFBs requires its ChT domain and methylated histone binding

TopoIIα and TopoIIβ are most divergent in their C-terminal domains (CTDs). The TopoIIα CTD (αCTD) has been found to interact with histones, preferentially with histone H3 tri-methylated at Lysine 27, and recent structural evidence indicates that the αCTD is positioned favorably for interaction with nucleosomes (Vanden Broeck et al., 2021). Based on these findings, we probed whether the difference in the abilities of TopoIIα and TopoIIβ to prevent UFBs arose from differing abilities to bind nucleosomes. To test this, we designed mono-nucleosome pull down assays. Recombinant S-tagged αCTD and TopoIIβ-CTD (βCTD) were used as bait to bind mononucleosomes prepared from salt extracted chromatin, digested by MNase (Supplemental fig. 3). Nucleosomes bound to bait proteins were analyzed by western blot with antibodies against histone species. Both CTD fragments bound to nucleosomes, indicated by the presence of H4 and H3 in the precipitates. However, only the αCTD could bind H3 with methylated lysine 27 (H3K27me3 and H3K27me2) (Fig. 2 A). This revealed that even though both these isoforms bind to nucleosomes, TopoIIα alone can bind specifically to nucleosomes containing H3K27me2/3. The results demonstrate that the previously established ability of the αChT domain to interact with H3K27me3 tail peptides (Lane et al., 2013), also applies to native nucleosomal H3K27me3. The αChT contains three aromatic amino acids that could contribute to recognition of trimethylated lysine, similar to the mechanism employed by HP1 and PRC2 where an aromatic “cage” surrounds a tri-methylated lysine residue (Jacobs and Khorasanizadeh, 2002; Min et al., 2003; Nielsen et al., 2002). Further, αChT and the corresponding region in TopoIIβ have differences in their aromatic amino acid composition (Fig. 2 B), suggesting that the difference in abilities of the two isoforms to bind methylated histone H3 and to completely resolve catenations, could arise from the properties of the αChT.

**Figure 2.**
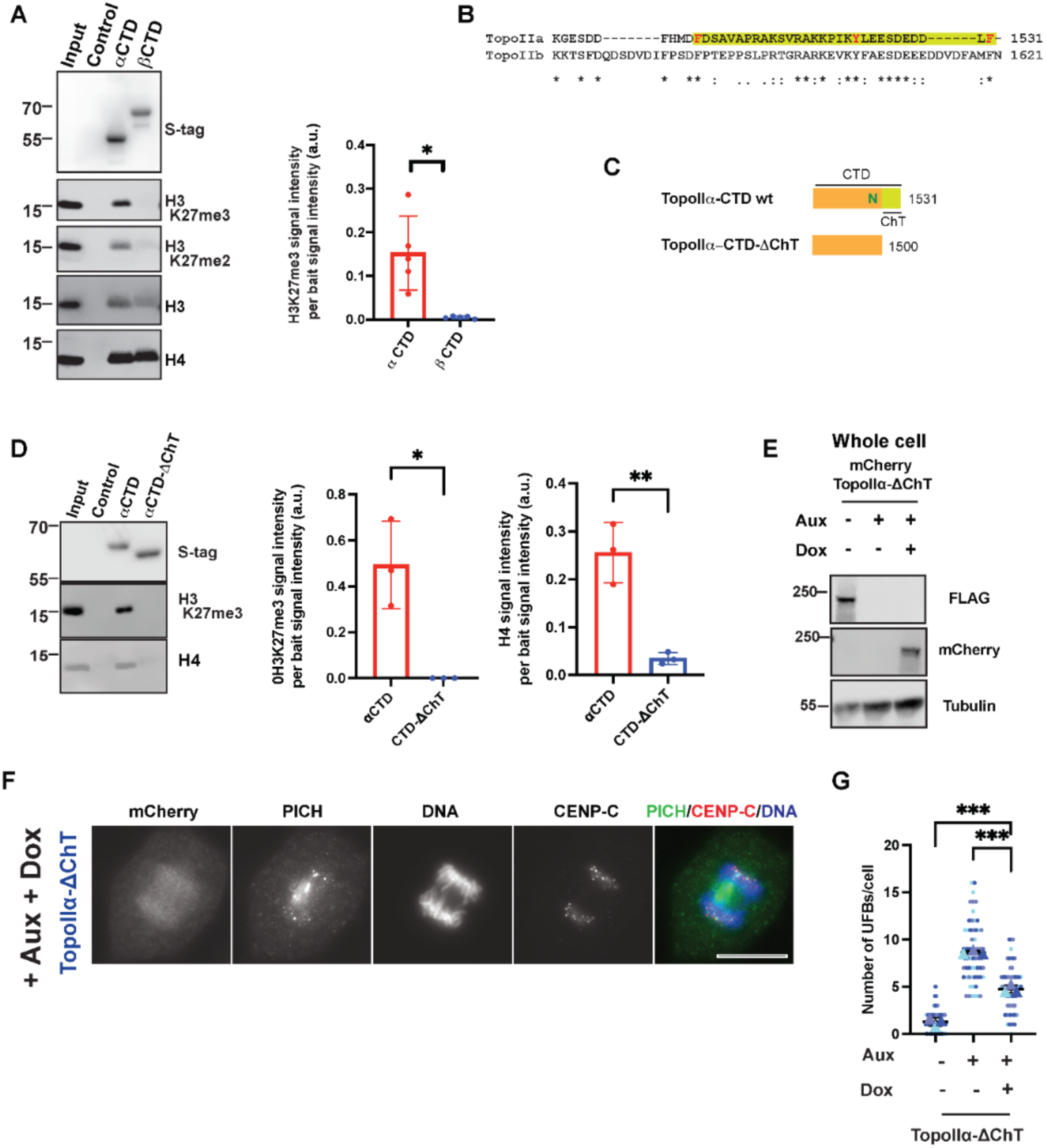
Complete UFB prevention by TopoIIα’s requires its ChT domain: **A)** Precipitated fractions of mono-nucleosome pull down assay with TopoIIα-CTD (αCTD) and TopoIIβ-CTD (βTD) were probed with indicated antibodies. H3K27me3 signal intensity per bait signal intensity (a.u.) was quantified over N=5 experiments. P-value indicates two-tailed unpaired samples t-test. Error bars indicate SD. *: p<0.033. **B)** Primary structure representation of the TopoIIα-ChT domain (highlighted in yellow) compared to the corresponding region in TopoIIβ. **C)** Representation of TopoIIα-CTD wt and its ChT domain-truncated mutant (highlighted in yellow) (TopoIIα-CTD-ΔChT). **D)** Western blot from mono-nucleosome pull down assay comparing αCTD wt and αCTD-ΔChT (left) with indicated antibodies. H3K27me3 and H4 signal intensities per bait intensity (a.u.) were quantified N=3 experiments (right). P value indicates two-tailed unpaired samples t-test. Error bars indicate SD. *: p<0.033, **: p<0.002. **E)** Western blot indicating TopoIIα-replacement with mCherry tagged TopoIIα–ΔChT (ΔChT) in cells. **F)** UFB assay with TopoIIα–ΔChT mutant replacement by staining with indicated antibodies. Bars, 10 μm. **G)** Superplots showing quantification of the number of UFBs/cell (from >60 cells counted over three independent experiments) for TopoIIα-ChT replaced cells. P-value indicates one-way ANOVA analysis followed by Tukey multicomparison correction for the means. Horizontal bars indicate mean and error bars indicate SD calculated for the means across the three independent experiments. ***: p<0.001.

To explore this idea, we performed mono-nucleosome pull down assays with a mutant αCTD that lacks the αChT domain (αCTD-ΔChT) (Fig. 2 C). We found that the αChT is required for this αCTD-specific binding to methylated H3 containing nucleosomes, because αCTD-ΔChT showed no detectable H3K27me3 precipitation. αCTD-ΔChT also showed a significantly reduced ability to precipitate histone H4 (Fig. 2 D). We then tested if this loss of binding to methylated H3 nucleosomes correlates with the inability to completely prevent UFBs. The depletion of endogenous TopoIIα and replacement with TopoIIα–ΔChT was tested using western blot (Fig 2 E). Deletion of the αChT (mCherry-TopoIIα–ΔChT) led to a chromosomal localization defect consistent with that previously reported (Lane et al., 2013), having reduced propensity for axial localization (Fig. 2F) as compared to wt (Fig.1 H). This mutant could decrease UFBs, but the rescue remained incomplete (Fig. 2 G). The incomplete rescue of UFBs by TopoIIα–ΔChT is comparable to that when TopoIIα was replaced with TopoIIβ wt (Fig. 1 I – in shades of red), suggesting that the divergent ChT domains could account for the requirement of TopoIIα, not TopoIIβ, for resolving a subset of catenated loci. The requirement of αChT for the complete resolution of catenations is further supported by data from experiments utilizing mutants in which the αChT and the region of TopoIIβ corresponding to αChT were swapped (Supplemental fig. 4). Mono-nucleosome pull down assays showed both αCTD and βCTD-αChT precipitated H3K27me3-containing nucleosomes (Supplemental fig. 4 B; lanes 2 and 5), but αCTD-ßChT did not. Thus, the αChT governs specific binding of the αCTD to nucleosomes containing H3K27me3. Quantification of UFBs in TopoIIα-replaced cells with expression of mCherry-TopoIIα-βChT or mCherry-TopoIIβ-αChT further support the requirement of αChT for complete UFB rescue (Supplemental fig. 4 D and E). These results from the “ChT-swapped” mutants support the requirement of the αChT in resolving a TopoIIα-dependent subset of catenated loci and the potential contribution of αChT/H3K27me3 binding for complete resolution of catenations.

### Disruption of αChT/H3K27me3 interaction increases UFBs

Aromatic amino acids can create a binding pocket for a tri-methylated lysine, as occurs in well-established trimethylated H3 binding proteins, HP1 for H3K9me3 and PRC2 for H3K27me3 (Jacobs and Khorasanizadeh, 2002; Min et al., 2003; Nielsen et al., 2002). Upon comparing the primary structures of the two isoforms among vertebrates, we noticed that the region corresponding to αChT is more divergent than the corresponding βChT region (Supplemental fig. 5 A and B) and the aromatic amino acid composition of αChT is not fully conserved among vertebrates. However, when we compared the primary sequence of αChT among primates there is a very high degree of conservation in this region (Supplemental fig. 5 C). We therefore focused on three aromatic residues (F1502, Y1521, F1531) within the αChT and investigated their specific role in H3K27me3 interaction and prevention of UFBs by mutating them to alanine (Fig. 3 A). In the mono-nucleosome pull down assay, all three mutants had reduced nucleosome binding, indicated by reduced H4 and H3 precipitation compared to αCTD wt (Fig. 3 B). Among them, Y1521A showed the least binding, a deficit that was similar to αCTD-ΔChT (Fig. 2 D). H3K27me3 binding was almost abolished in both F1502A and Y1521A. The F1531A precipitated significantly less H3K27me3-containing nucleosomes than αCTD wt but did precipitate significantly more than both F1502A and Y1521A (Fig. 3 B right panel).

**Figure 3.**
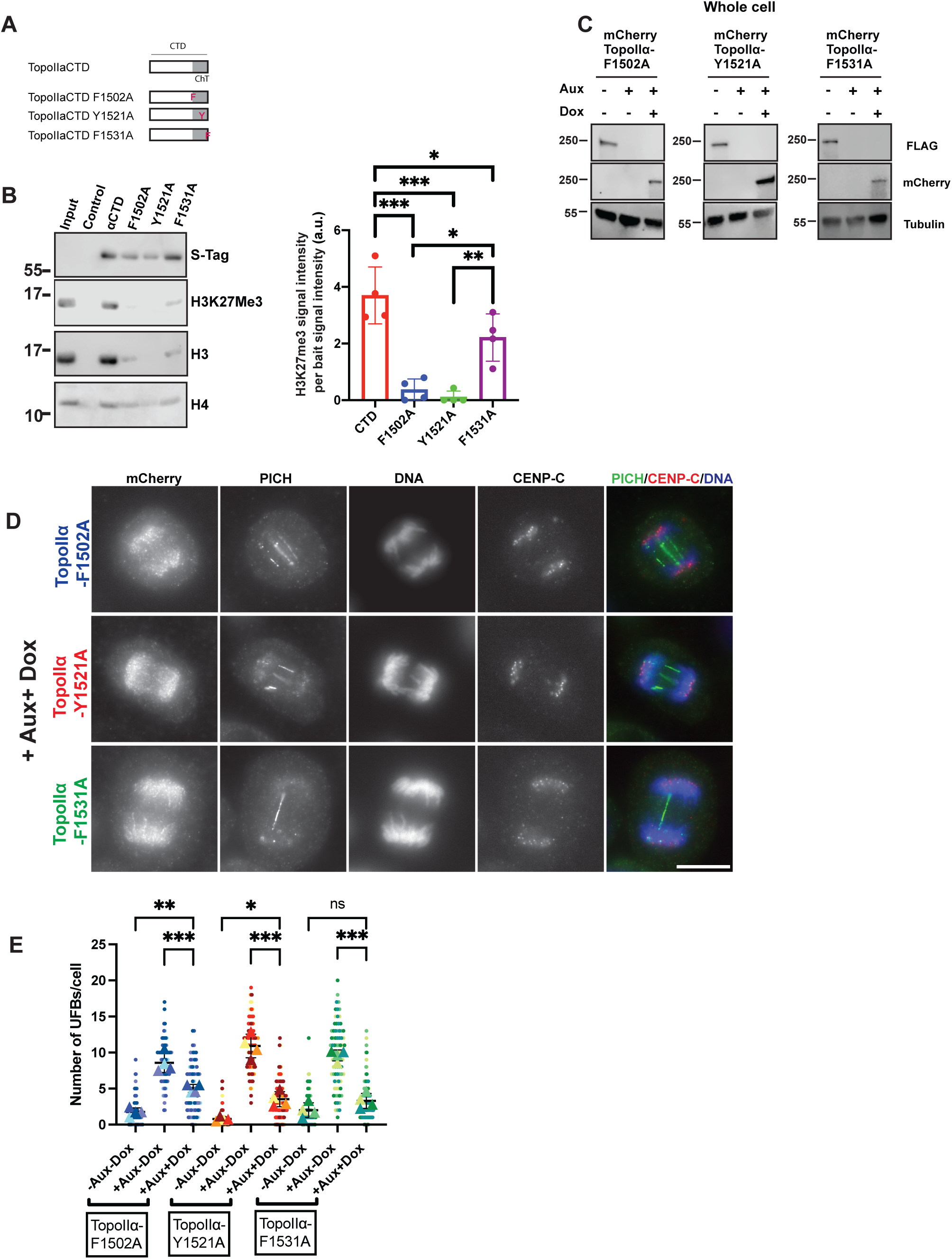
H3K27me3 binding and UFB assays indicate that aromatic amino acids in TopoIIα-ChT play differential roles. **A)** Representation of TopoIIα point mutants. Each of the aromatic residues (refer to Fig. 2 B) Phel502, Tyrl52l and Phel53l were replaced with Ala (represented as F1502A, Y1521A and F1531A respectively). **B)** Binding of each TopoIIα point mutant CTD to H3K27me3 containing chromatin indicated by mono-nucleosome pull down assays (left). H3K27me3 signal intensity per bait signal intensity (a.u.) bound by the various TopoIIα point mutants is summarized from N=4 experiments (right). P-value indicates one-way ANOVA analysis followed by Tukey multicomparison correction. Error bars indicate SD. *: p<0.033, **: p<0.002, ***: p<0.001. **C)** Western blot showing replacement with the TopoIIα point mutants. **D)** Representative images of UFB assay with TopoIIα point mutants’ replacement by staining with antibodies against indicated targets. Bars, 10 μm. **E)** Superplots showing quantification of the number of UFBs/cell (from >60 cells counted over three independent experiments) for replacement with TopoIIα point mutants. P-value indicates one-way ANOVA analysis followed by Tukey multicomparison correction for the means. Horizontal bars indicate mean and error bars indicate SD calculated for the means across the four independent experiments. ns: not statistically significant, *: p<0.033, **: p<0.002, ***: p<0.001.

To examine the relationship between H3K27me3 binding and prevention of UFBs, we created cell lines with endogenous TopoIIα replaced with mutant TopoIIα carrying the same aromatic residue mutations. The replacements were confirmed, as for the other cell lines (Fig. 3 C) and UFB assays performed (Fig. 3 D). Quantification of UFBs in each mutant revealed data consistent with a significant contribution of H3K27me3-containing nucleosome binding for complete prevention of UFBs. Both TopoIIα-F1502A and TopoIIα-Y1521A showed incomplete prevention of UFBs, similar to TopoIIβ and TopoIIα–ΔChT with significantly increased numbers of UFBs compared to controls. Intriguingly, the TopoIIα-F1531A did not have a statistically significant increase in the number of UFBs compared to control (Fig. 3 E). The F1531A also retained weak but significant affinity to H3K27me3-containing mono-nucleosomes, unlike the other two mutants. The results from the three mutants are consistent with H3K27me3 binding of TopoIIα, governed by the αChT, playing an important role in ensuring complete resolution of catenations to prevent UFBs.

### Methylated histone binding dependent on the αChT regulates association of TopoII with mitotic chromosomes

Loss of the αChT was shown to decrease the association of TopoIIα with chromosomes in live mitotic cells (Lane et al., 2013). To ask if the aromatic αChT residues that are required for interaction with H3K27me3 dictate this association of TopoIIα with chromosomes in live mitotic cells, we imaged the mutants utilizing the mCherry fused to these proteins. The entire cell volume of nocodazole arrested cells was captured at 0.2μm intervals and mCherry-TopoII signal intensity was quantified in the projected images (Fig. 4A). Quantification of chromosomal mCherry-TopoIIα signals revealed a decrease in the H3K27me3 binding deficient mutants, consistent with previous observations with the ΔChT mutant (Lane et al., 2013) (Fig. 4B). Among the aromatic amino acid mutants, F1502A and Y1521A had decreased TopoII association with chromosomes. However, F1531A, which partially retains H3K27me3 binding ability, and was able to rescue the UFB phenotype, did not show a reduction in chromosome association compared to wt. Therefore, we observed that a correlation exists between the ability to bind H3K27me3, the association with mitotic chromosomes, and the ability to prevent UFBs. The signal intensities of each protein were also measured in interphase cells in which they were expressed and were found to be consistent with each other (Fig 4 C). This indicates that the differences in their mitotic chromosomal signal intensity originates from their differential abilities to associate with mitotic chromosomes and not from variations in their expression levels.

**Figure 4.**
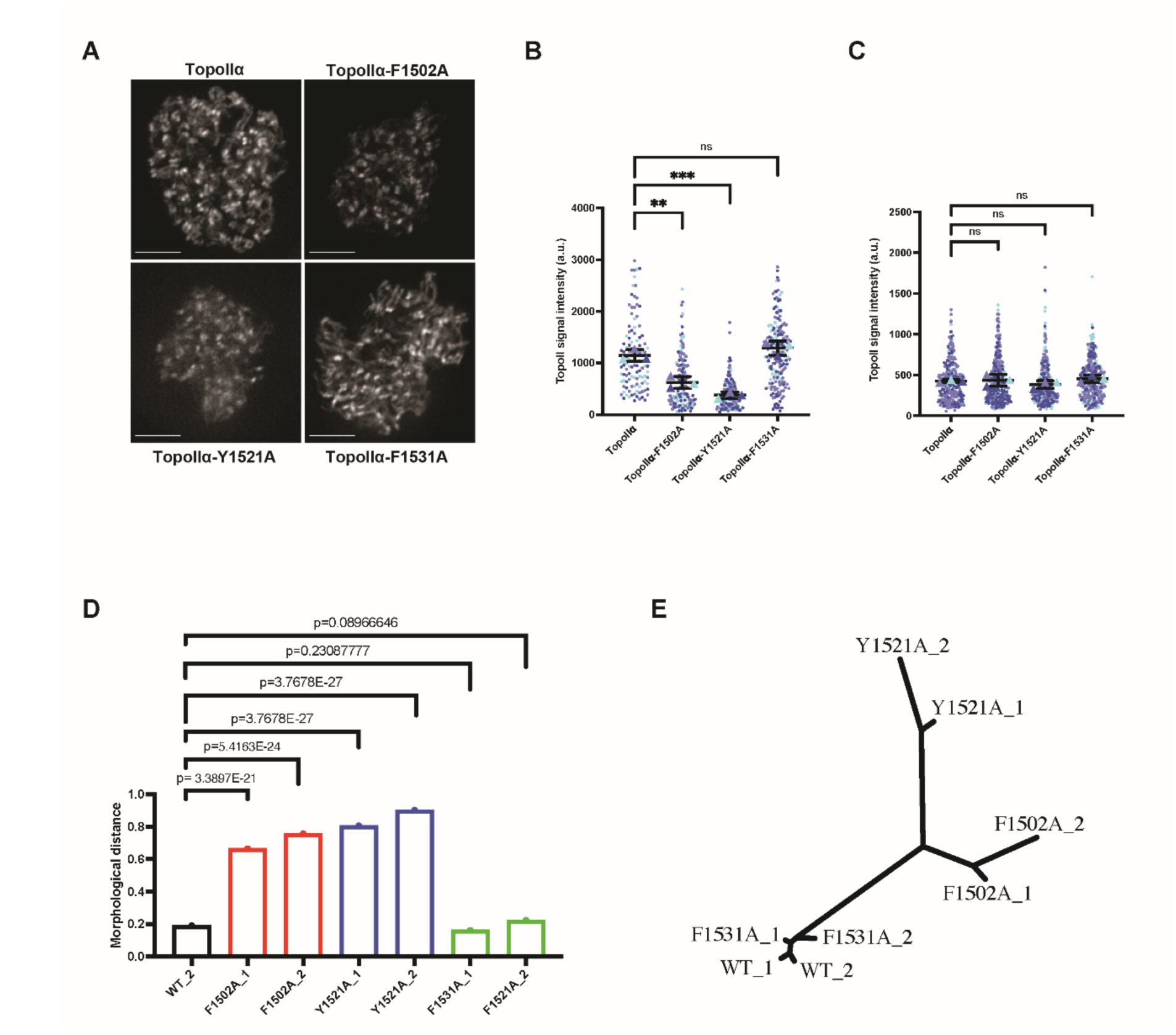
TopoIIα-ChT dependent methylated histone binding regulates association of TopoII with mitotic chromosomes. **A)** Projected images of mCherry-TopoII in live mitotic cells. Bars, 5 μm. **B)** Superplots showing quantification of mCherry-TopoII signal intensity on mitotic chromosomes in live cells (from >100 cells counted over three independent experiments). P-values indicate one-way ANOVA analysis followed by Tukey multicomparison correction. Horizontal bars indicate means and error bars indicate SD calculated for the means across the three independent experiments. ns: not statistically significant; **: p<0.002, ***: p<0.001. **C)** Superplots showing quantification of mCherry-TopoII signal intensity on interphase nucleus in live cells (from >150 cells counted over three independent experiments). P-values indicate one-way ANOVA analysis followed by Tukey multicomparison correction. Horizontal bars (black) indicate means and error bars (colors) indicate SD calculated for the means across the three independent experiments. ns: not statistically significant. **D)** Morphological distances of the indicated types of TopoIIα from wt (WT_1). Each group was randomly divided into two subgroups to monitor reproducibility and reliability of the analysis. A larger distance indicates a greater morphological dissimilarity in an image feature space. While F1502A and Y1521A were distinct from wt TopoIIα, F1531A were similar to it. **E)** Phylogenetic tree representing the morphological relationships among the indicated TopoIIα. The tree is based on the values obtained from the wndchrm analysis in Fig. 4 D.

In addition to the differences in signal intensities, there were variations in the localization patterns among the wt and TopoIIα mutants, particularly in the CEN and chromosome axial distributions (Fig 4 A). To gain an unbiased evaluation of their general morphologies, we applied the mCherry-TopoIIα images to a machine leaning algorithm, wndchrm (weighted neighbor distance using a compound hierarchy of algorithms representing morphology) (Shamir et al., 2008). Initial image-titration analysis showed that the classification accuracy was high at 88% with 10 training images of the wt and the mutant TopoIIα (Y1521A) (supplemental fig. 6). We hence collected 30 images from each TopoIIα cell line for the analysis, which were randomly subdivided into two sub-groups: WT_1 and WT_2, for example. The resultant eight subgroups were subjected to wndchrm analysis to measure differences by computing the morphological distances (Fig. 4D). As expected, distances between the two subgroups from the same protein were small, confirming the accuracy of the analysis. The morphological distances of F1502A and Y1521A from wt were large, whereas that of F1531A was far less. Further, the morphological similarity and dissimilarity was visualized as a phylogenetic tree (Fig. 4E), which showed that F1502A and Y1521A were morphologically distant from wt, as well as from each other. These results indicate that methylated histone binding of αChT has an additional role in facilitating association of TopoIIα with mitotic chromosomes.

### Reducing H3K27me3 via EZH2 methyltransferase inhibition induces chromosome segregation defects and hinders proper TopoIIα localization on mitotic CENs

Thus far, our results suggest that the αChT/H3K27me3 interaction contributes to TopoIIα’s function in completely resolving the catenated genome and in TopoIIα localization to mitotic chromosomes. In order to examine if H3K27me3 is indeed a key contributor to these functions, we utilized a well-established H3K27 tri-methylase inhibitor, GSK-343. Previous studies used a range of concentrations (0.110 μM) and treatment times (1-7 days) based on the cell lines and experimental protocols utilized (Mohammad et al., 2017; Verma et al., 2012). To determine the effect of the inhibitor under our experimental conditions, we treated cells with varying concentrations of GSK-343 (2-6μM) after thymidine release and collected the mitotic chromosomes after 8.5 hours (Fig. 5 A). Western blot of these chromosomes indicated a reduction in H3K27me3 with GSK-343 treatment at all concentrations. The H3K27me3 signal reduction demonstrated concentration dependency that plateaued at 4 μM GSK-343 (Fig. 5 B). This data suggests that there exists a subset of H3K27 that undergoes tri-methylation between S-phase and mitosis, perhaps consistent with studies that found a decrease in H3K27 acetylation as cells progress into mitosis (Kang et al., 2020). We found that GSK-343 did not affect H3K9me3 levels under the same conditions (Fig. 5 C). Following these observations of the effect of GSK-343 on H3K27me3 on mitotic chromosomes, we examined the requirement of H3K27me3 in αChT-mediated functions using our established treatment conditions and 4 μM GSK-343 (Fig. 5 A).

**Figure 5.**
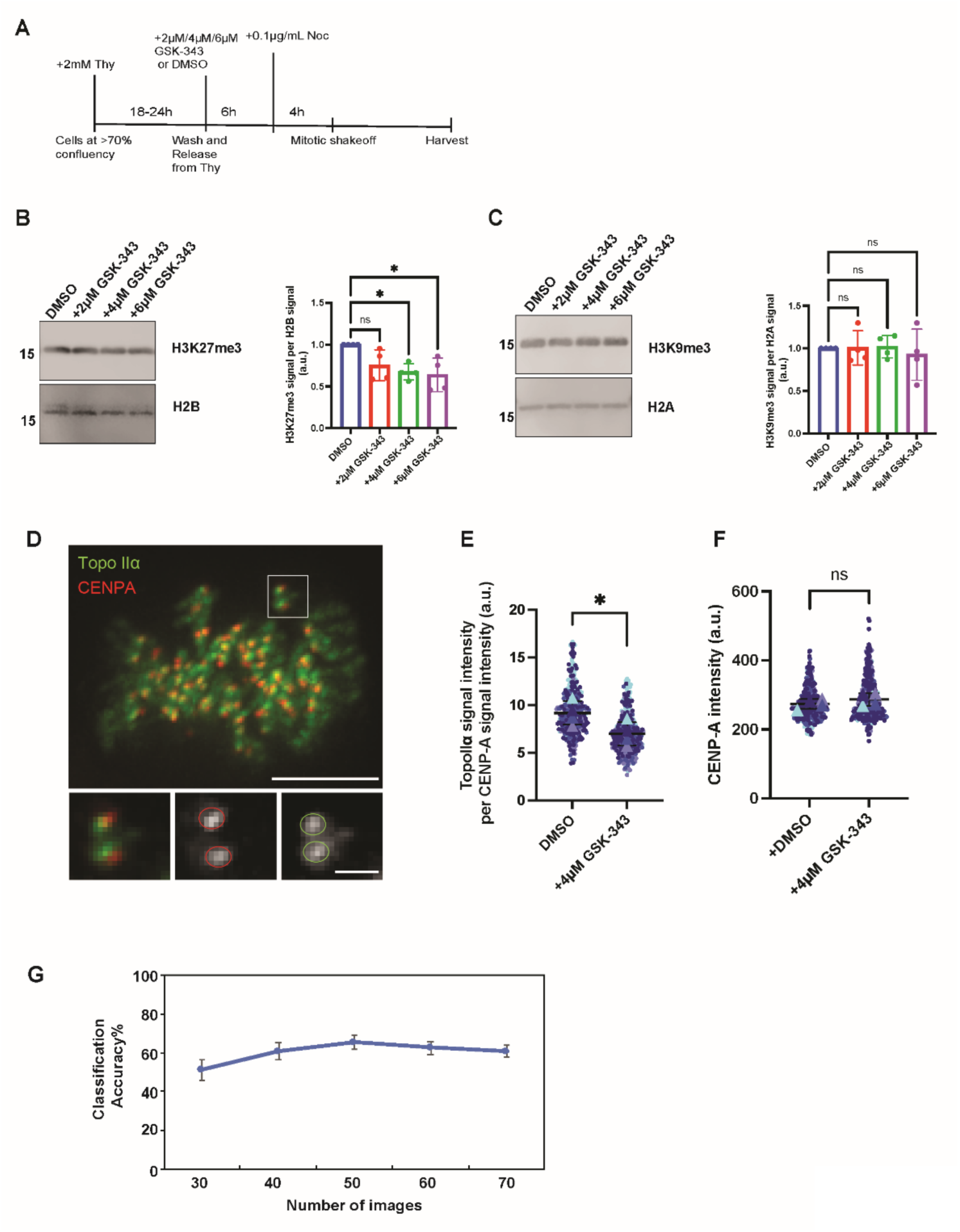
H3K27me3 inhibition by GSK-343 results in reduction of TopoIIα CEN signal. **A)** Schematic of synchronization method and harvesting of chromosomes with GSK-343 (2 μM/4 μM/6 μM) or DMSO (control) treatments. **B/C)** Western blot of mitotic chromosomes with GSK-343 treatment for H3K27me3 amount (B) and H3K9me3 amount (C) on chromosomes. Representative gel images are shown in the left panels and quantification of signal intensities are shown in the right panels. P-values indicate one-way ANOVA analysis followed by Tukey multicomparison correction. Horizontal bars indicate means and error bars indicate SD calculated for the means across the four independent experiments. ns: not statistically significant; *: p<0.033. **D)** Representative images of live cells with CEN TopoIIα and CENP-A (CEN marker) used for quantification of CEN TopoIIα signal. Bars, 5μm (larger merged image), 1 μm (magnified images). **E)** Superplots for quantification of TopoIIα signal intensity at CENs against the corresponding CENP-A signal intensity for both DMSO and GSK-343 treatment. The data is from 366 chromosomes for DMSO condition and 413 chromosomes for GSK-343 treated condition counted over 3 independent experiments as represented in the plot. P value indicates two-tailed unpaired samples t-test. Error bars indicate SD. *: p<0.033. **F)** Superplots for quantification of CENP-A signal intensity from CENs used for quantification in Fig. 5 E. P value indicates two-tailed unpaired samples t-test. Error bars indicate SD. ns: not statistically significant **G)** Classification accuracies measured by wndchrm to distinguish between TopoIIα in the DMSO and GSK-343 treated cells. Approximately 50% accuracies were obtained regardless of the training numbers (30-70 images), indicating that TopoIIα localization did not change globally by the GSK-343 treatment.

First, we tested the effect of GSK-343 on TopoIIα association with mitotic chromosomes using a cell line established with endogenous TopoIIα fused with mNeon as well as endogenous CENP-A fused with miRFP (supplemental fig. 7). With this line, we were able to perform live imaging of mitotic chromosomal TopoIIα as we were able to do with the mutants in fig. 4. CENP-A-miRFP signals were used as a guide for determining the CEN regions at which TopoIIα is known to localize by forming foci. Images from Pro-metaphase cells with distinguishable chromosomes, as shown in an example (Fig. 5 D), were chosen for quantification of CEN TopoIIα foci. The results showed a significant reduction in CEN TopoIIα signal intensity after GSK-343 treatment (Fig. 5 E) that was normalized against CENP-A signal intensity (Fig. 5 F), which showed no significant changes with GSK-343. Because mutants of the αChT showed defects in chromosomal localization, we probed into whether the global localization of TopoIIα changes upon GSK-343 treatment. We measured classification accuracies for TopoIIα in the control and the drug treated cells, using the wndchrm algorithm (Shamir et al., 2008). For this analysis, we used images of TopoIIα distributed on the entire chromosomes. We found that regardless of the increasing numbers of the training images (from 30 to70 images), classification accuracies stayed ~50% (Fig. 5 G). This indicates differences in global TopoIIα localization between control and GSK-343-treated cells are subtle. This may be consistent with the limited reduction of H3K27me3 after GSK-343 treatment (Fig. 5 A-C), probably because the treatment was for a short period and at specific cell cycle stages. Only a subpopulation of H3K27me3 may be established from S-phase to mitosis, which contributes to CEN TopoIIα localization. Global chromosomal localization of TopoIIα may be insensitive to this treatment regimen.

### H3K27me3 is required for proper chromosome segregation and complete prevention of UFBs

To ask if H3K27me3 contributes to αChT-dependent mitotic functions, we quantified chromosome bridge formation after GSK-343 treatment. Utilizing the mNeon-TopoIIα line and live imaging, we counted cells with anaphase chromosome bridges with 4 μM GSK-343 treatment (and control cells with DMSO) (Fig. 6 A).

**Figure 6.**
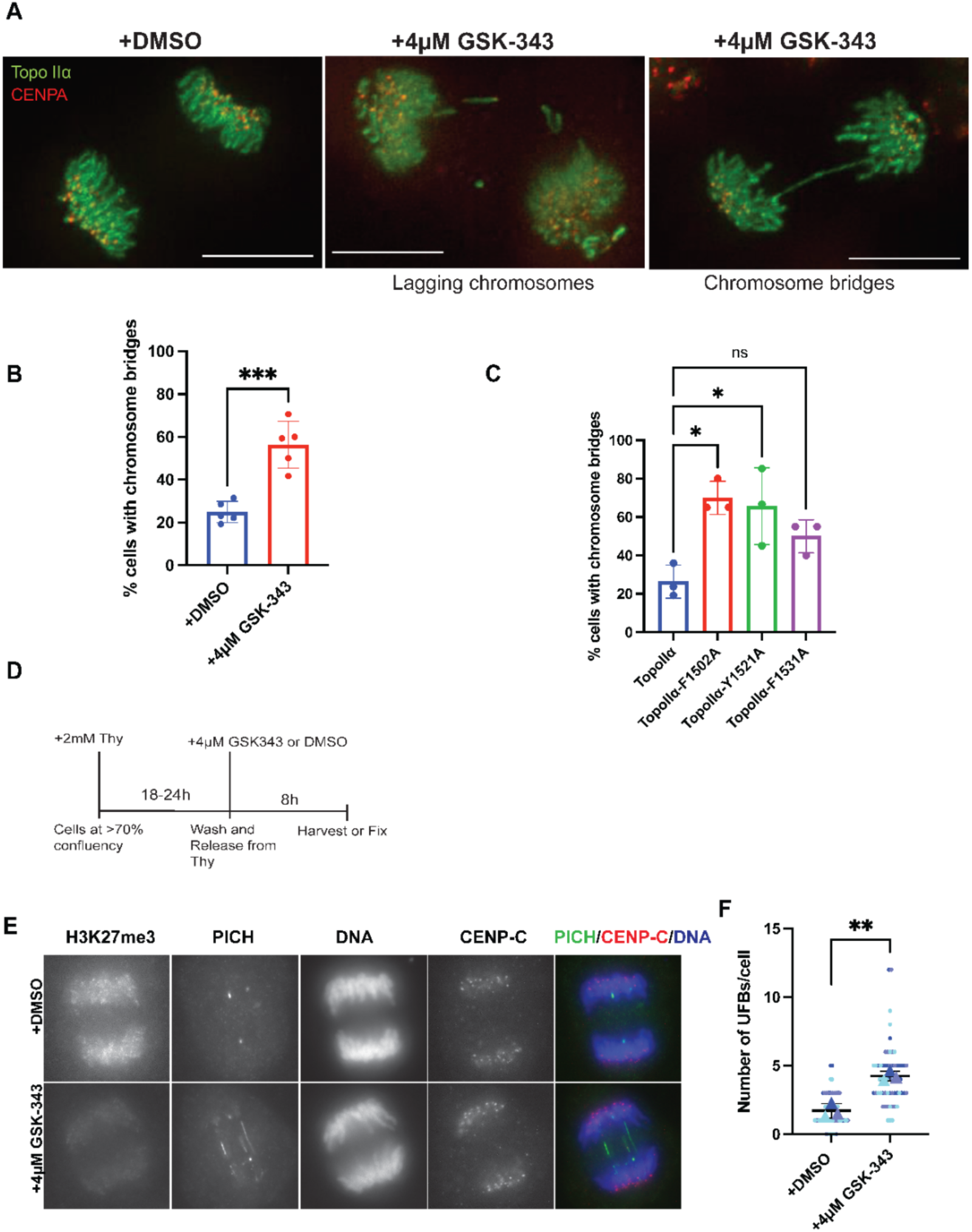
H3K27me3 inhibition results in increased chromosome bridges and UFBs in anaphase cells. **A)** Projected images of mNeon-TopoIIα, miRFP-CENP-A in live mitotic cells showing chromosome bridges and/or lagging chromosomes. Bars, 5 μm. **B)** Quantification of cells with chromosomes bridges with GSK-343 treatment from N=5 experiments, with 154 (DMSO) and 174 (GSK-343) total cells counted. P value indicates two-tailed unpaired samples t-test. Error bars indicate SD. ***: p<0.001. **C)** Quantification of cells with chromosome bridges in cells replaced with ChT point mutants from N=3 experiments with at least 20 cells counted for each condition. P-values indicate one-way ANOVA analysis followed by Tukey multicomparison correction. Horizontal bars indicate means and error bars indicate SD. ns: not statistically significant; *: p<0.033. **D)** Schematic of cell synchronization method for UFB assays with GSK-343 treatment. **E)** Representative images of UFB assays with 4 μM GSK-343 treatment. Cells were stained with indicated antibodies. **E)** Superplots showing quantification of the number of UFBs/cell (from >60 cells counted over three independent experiments) for UFB assays with 4 μM GSK-343 treatment. P-value indicates one-way ANOVA analysis followed by Tukey multicomparison correction for the means. Horizontal bars indicate mean and error bars indicate SD calculated for the means across the three independent experiments. **: p<0.002.

With inhibition of H3K27me3, on average half of the population displayed chromosome bridges in comparison to an average of 20% of control cells (Fig. 6 B). This increase in cells with chromosome bridges with GSK-343 treatment is consistent with those observed in the αChT-mutants, measured from fixed cell images (Fig. 6 C). We also observed lagging chromosomes (Fig. 6 A, center) upon treatment with GSK-343. We next asked if H3K27me3 is directly linked to αChT-dependent prevention of UFBs by quantifying UFBs after GSK-343 treatment. Cells were treated with GSK-343 after thymidine release (Fig. 6 D) and subjected to UFB analysis by PICH staining. H3K27me3 was visualized by anti-H3K27me3 staining. With 4 μM GSK-343, there was a decrease in H3K27me3 on anaphase chromosomes and an increase in the number of PICH-coated UFBs (Fig. 6 E). We also examined the effect of 2 μM and 6 μM GSK-343 (supplemental fig. 8) and found that there was a dosage dependent increase in the number of UFBs, with saturation at 4 μM GSK-343. This increase in UFBs/cell was statistically significant when compared to the control condition (Fig. 6 F) and was closely comparable to the average UFBs/cell in the TopoIIα–ΔChT. This demonstrates that the proper establishment of H3K27me3, presumably on CEN nucleosomes between S-phase and mitosis, is required for the complete resolution of sister chromatid catenations.

To determine if the increased UFBs after GSK-343 treatment are due to a functional relationship between the αChT and H3K27me3, we examined the effect of GSK-343 in the αChT mutants that cannot bind to H3K27me3 (i.e., F1502A and Y1521A). Cells were treated with Aux and Dox as well as 4 μM GSK-343 following the release of cells from thymidine synchrony (Fig. 7 A). This revealed a significant increase in UFBs after GSK-343 treatment in mCherry-TopoIIα wt replaced cells (Fig 7B and 7C), similar to that observed in Fig. 6 E and F. Importantly, in the mutants that have an H3K27me3 binding deficiency– TopoIIα-F1502A (Fig 7D and 7E) and TopoIIα-Y1521A (Fig 7F and 7G) - there was no significant difference between control cells and the GSK-343 treated cells. This reveals that there is no synergistic effect of H3K27me3 reduction and αChT mutations that decrease binding to H3K27me3. The same outcome was observed with TopoIIα-βChT, where H3K27me3 binding is compromised, but there was a synergy between GSK-343 treatment and TopoIIα-F1531A, that retains some H3K27me binding ability (supplemental fig. 9). This data reinforces the evidence that binding of the αChT to H3K27me3-containing nucleosomes is a critical mechanism required for complete resolution of sister chromatids for faithful mitotic segregation.

**Figure7.**
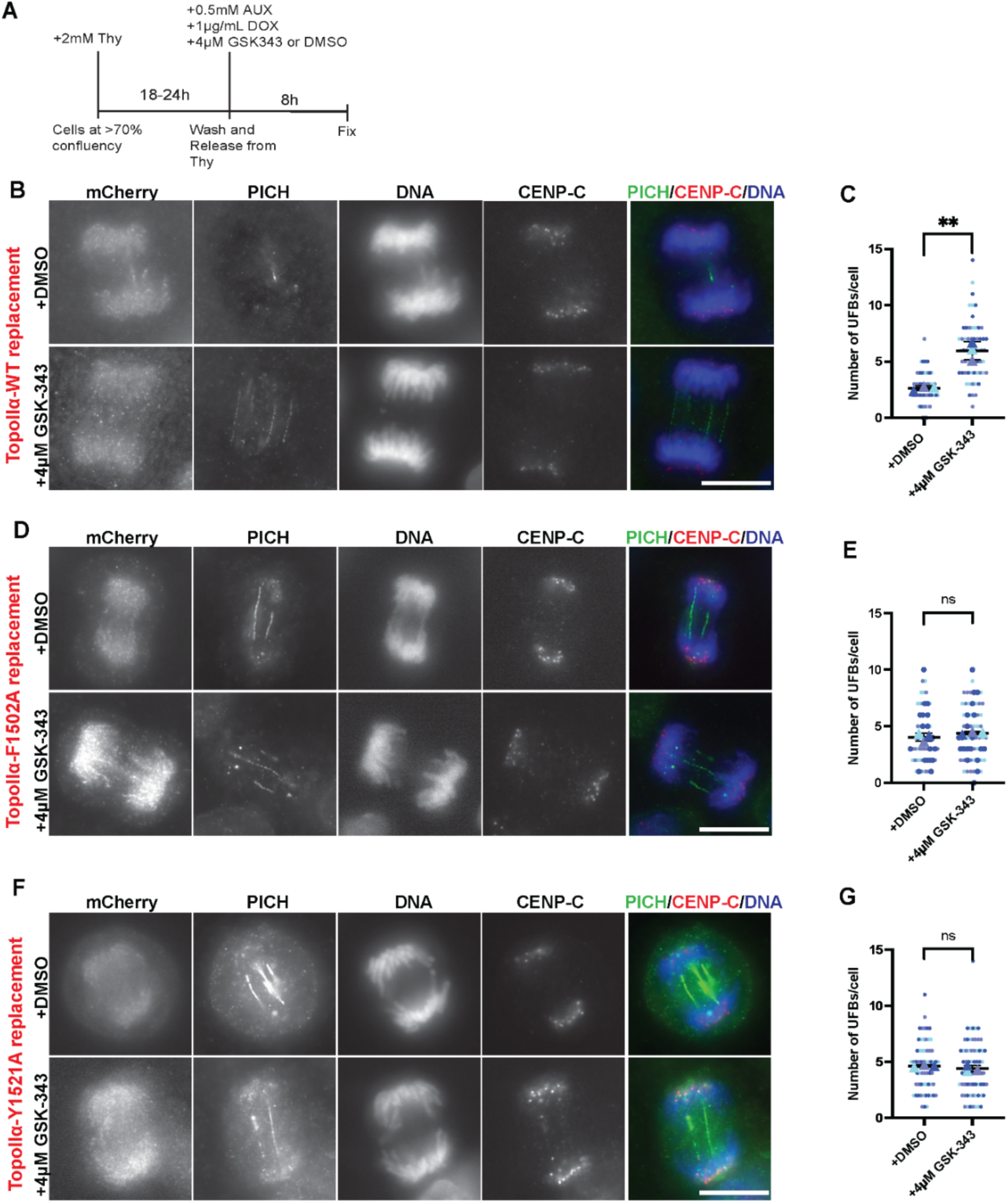
H3K27me3 inhibition shows no synergistic increase in UFBs in cells replaced with H3K27me3 binding deficient TopoIIα mutants. **A-** Schematic of cell synchronization method for UFB assays with endogenous TopoIIα-replacement to mCherry-TopoIIα wt/mutants along with the addition of 4 μM GSK-343 (or DMSO in control cells). **B/D/F**-Representative images of UFB assay in TopoIIα-replaced cells with mCherry-TopoIIα wt (B), mCherry-TopoIIα F1502A (D), and mCherry-TopoIIα Y1521A (F). Cell with indicated treatment, DMSO or 4 μM GSK-343 were stained with indicated antibodies for UFB measurement. Bars, 10 μm. **C/E/G-** Superplots showing quantification of the number of UFBs/cell (from >60 cells counted over three independent experiments) for TopoIIα wt replacement (C), TopoIIα F1502A replacement (E), and mCherry-TopoIIα Yl52lA-replacement (G). P-value indicates one-way ANOVA analysis followed by Tukey multicomparison correction. Horizontal bars indicate mean and error bars indicate SD calculated for the means across the three independent experiments. ns: not statistically significant, **: p<0.002.

## Discussion

### Role of TopoIIα-ChT in resolving catenated centromeres

Altogether, our results throw new light on one of the novel mechanisms through which TopoIIα can detect catenated genomic DNA to facilitate efficient and complete separation of the sister chromatids in anaphase. UFBs seen in early to mid-anaphase are known to be catenated DNA molecules that link the sister CENs (Chan et al., 2007; Ke et al., 2011). While bulk decatenation of the genome may not be highly regulated, the ability of TopoIIα to specifically localize to a small number of persistent catenations appears to be reliant on a highly targeted mechanism to achieve the necessary precision for faithful chromosome segregation within a short timeframe. The data presented here provide evidence that methylated histone binding, specifically H3K27me3, by the αChT is critical for TopoIIα-specific recognition of the last remaining catenated CEN loci. This mechanism of chromatin recognition can account for the distinct requirement of the α-isoform vs. the ß-isoform for the prevention of UFBs. The data are consistent with a model where a proportion of the catenated genome can be recognized by either isoform, because there was significant rescue of UFBs by ectopic over-expression of TopoIIβ in TopoIIα-depleted cells (Fig 1 G). It is noteworthy, however, that the association of TopoIIβ with mitotic chromatin under normal conditions is very low compared with TopoIIα (Fig 1D). Thus, the rescue of UFBs by ectopic over-expression of TopoIIβ may over-estimate any biologically relevant contribution of TopoIIβ to chromosome segregation. Furthermore, our data clearly revealed that there are specific catenated CEN loci that cannot be resolved efficiently by TopoIIβ, and these loci require a TopoII with an αChT that can interact with H3K27me3 modified nucleosomes. The αChT and βChT domains are strikingly distinct in their abilities to interact with H3K27me3-containing chromatin. Indeed, comparing the binding ability to H3K27me3 and prevention of UFBs in all the mutants tested, the results strongly support the conclusion that methylated histone binding is required for complete resolution of αChT-dependent catenated CENs, i.e., the mutants that lack H3K27me3 binding ability cannot completely prevent UFBs (Fig. 3 and supplemental fig. 4). The contribution of H3K27me3 to the αChT-dependent function of TopoIIα is further supported by the results with GSK-343, a potent inhibitor of the H3K27 methyltransferase PRC2, which showed no synergistic increase in UFBs in the TopoIIα ChT mutants that lack H3K27me3 binding. Chromatin binding specificity via the αChT is also required for the proper localization of TopoIIα on mitotic chromosomes. Altogether, we propose that the αChT/H3K27me3 interaction is required for mitotic TopoIIα function for faithful genome segregation as summarized in Fig. 8.

**Figure 8.**
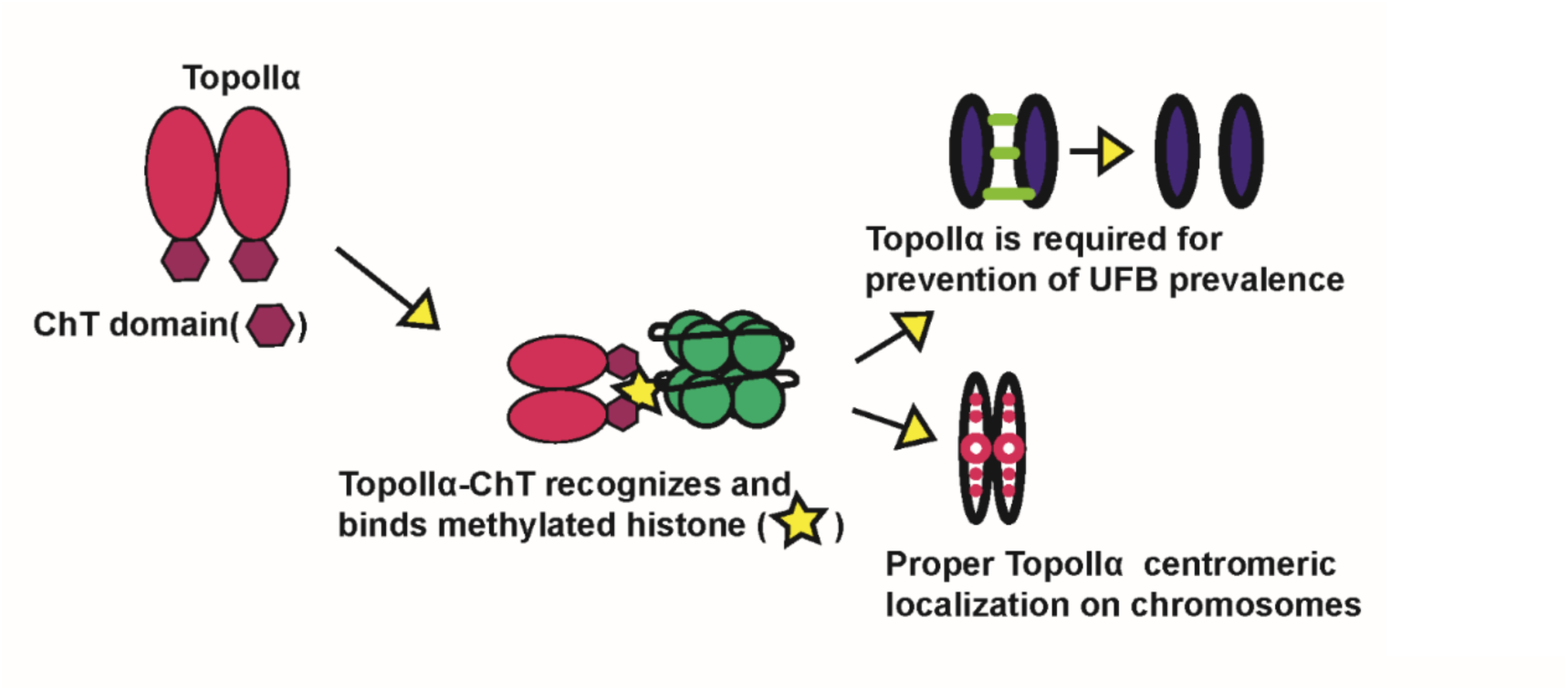
Mechanism of complete catenated genome resolution by TopoIIα. TopoIIα recognizes and binds methylated histones on chromatin, particularly H3K27me3, using its ChT domain. Through this mechanism, TopoIIα associates with chromatin and becomes targeted to catenated CEN loci. This binding therefore facilitates complete resolution of catenated loci, particularly CEN loci (for preventing UFBs) and chromosome bridges.

The evidence suggests that H3K27me3 is one of the critical histone modifications that marks catenated CEN loci. Indeed, it has been shown that H3K27me3 is specifically present at CENs in mitosis (Adriaens et al., 2016; Martins et al., 2016; Martins et al., 2020) and consistent with this, it is known that UFBs form when catenated CEN DNA molecules fail to be resolved before anaphase (Ke et al., 2011; Nielsen et al., 2015). Marking of catenated CEN loci with cues such as H3K27me3 is therefore a key mechanism that recruits TopoIIα to ensure resolution of the catenated CEN loci in a timely manner, before or during anaphase. Our results also suggest that the requirement of H3K27me3 for CEN TopoIIα localization (Fig. 5 D) and TopoIIα-dependent prevention of UFBs holds true for a subset of catenated CEN loci, but not for all. It will be imperative to further investigate if the H3K27me3 levels vary at each CEN and in-turn if the variation of H3K27me3 correlates with αChT-dependent TopoIIα association with and function on the chromosome. It has been proposed that TopoIIα preferably acts on positively supercoiled DNA over negatively supercoiled DNA, based on *in vitro* DNA relaxation assays. This substrate specificity requires the αChT region (Dickey and Osheroff, 2005). Therefore, it is plausible that catenated CEN loci that require the αChT for resolution, also possess positively supercoiled DNA that is needed for efficient decatenation by TopoIIα. The contribution/relationship of these two mechanisms for resolving catenated CEN loci will be important to investigate; for example, (1) If ablating histone modifications alters the topology of genomic DNA, and (2) If TopoII-mediated relaxation of supercoiled DNA is defective in the aromatic amino acid mutants.

### Role of αChT/H3K27me3 interaction in TopoIIα’s mitotic functions and regulation

We observed that TopoIIα-ChT aromatic amino acid mutants with reduced H3K27me3 binding ability had chromosome association defects (Fig. 4). This indicates the interaction with H3K27me3 via the αChT plays a critical role in TopoIIα chromosomal association. Further, we observed differences in the association ability of each mutant. F1502A and Y1521A displayed significant defects when compared to wt (Fig. 4 B). However, even among these two mutants we found differences in both nucleosome binding and localization on chromosomes (Figs. 3 B and 4 E). These observations call for further investigation into whether other histone modifications are also involved in controlling the proper association of TopoIIα with mitotic chromosomes. This possibility could also explain why the inhibition of H3K27me3 by GSK-343 did not result in global defects in chromosomal TopoIIα localization (Fig. 5 E), although it remains a possibility that more complete H3K27me3 reduction would be needed to greatly reduce the association of TopoIIα with chromosomes. Another finding was the increased number of chromosome bridges after treatment with GSK-343. Because of this, it may be that the αChT/H3K27me3 interaction contributes to additional aspects of chromosomal segregation, in addition to complete resolution of catenated CEN loci, since UFBs (by definition) are not associated with chromosomes bridges. In support of this possibility, we also found that αChT aromatic amino acid mutants (with lower H3K27me3 binding) had increased chromosome bridges, similar to GSK-343 treated cells. In line with this, a comprehensive identification of chromatin specifically bound to αChT will be critical to determine the whole picture of how the chromatin binding ability of TopoIIα functions in faithful genome transmission.

The aromatic amino acids that we focused on have varying degrees of conservation among vertebrates. Y1521, which we have found to be needed for both H3K27me3 binding and complete resolution of catenated CENs, is highly conserved among vertebrates (supplemental fig. 5 A). Although there is some discrepancy in the conservation of F1502 and F1531 among vertebrates, these residues are highly conserved among primates (supplemental fig. 5 C). Based on this observation, it will be important to probe into the potential of there being co-evolution between TopoIIα and the epigenetic cues that facilitate this novel TopoIIα regulatory mechanism.

### Complete resolution of the catenated genome via two distinct mechanisms?

Our results comparing TopoIIα and TopoIIβ indicate that the bulk of genome resolution can be performed by either of the isoforms and that the αChT domain is dispensable for this activity. It seems then, that the essential role of the αChT is to locate the last persistent catenations. The few CEN catenations that remain at the onset of anaphase contribute to sister chromatid cohesion in late metaphase (Diaz-Martinez et al., 2006). These are likely hard to identify with the required efficiency in the short time period from late metaphase to the end of anaphase. Rather than random attempts to locate these catenations, we propose that histone methylation, particularly H3K27me3, flags the loci and recruits TopoIIα via interaction with the αChT, thus facilitating faithful chromosome segregation. From a biological point of view, such a mechanism would be highly impactful because even single chromosomal aneuploidies have dramatic consequences including promotion of tumorigenesis.

## Materials and Methods

### DNA constructs, Recombinant proteins and antibodies Preparation

The parental line construct for OsTIR1 targeting and the AID-TopoIIα construct were created as described previously (Hassebroek et al., 2020). The mNeon fusion donor for endogenous TopoIIα was created by replacing AID sequence to mNeon sequence of TopoIIα N-terminal targeting donor plasmid. In this study, the donor plasmids for AID fusion to endogenous TopoIIβ at its N-terminal and miRFP670 fusion to endogenous CENP-A at its C-terminal were created. The homology arms for TopoIIβ N-terminal insertion and CENP-A C-terminal were amplified using primers listed in supplemental information (Table S1) from genomic DNA of DLD-1 cell. The amplified TopoIIβ homology arms were inserted into the plasmid by using PciI/SalI and SpeI/NotI sites in the 3x micro-AID/3x flag donor plasmid for described previously (Hassebroek et al., 2020). The amplified CENP-A homology arms were inserted into the plasmid by using PciI/SalI and SpeI/NotI sites in the C-terminal targeting donor plasmid carrying miRFP670 fused to puromycin resistant gene via T2A sequence as shown in supplemental fig S7. The guide RNA sequences for TopoIIβ N-terminal targeting and CENP-A C-terminal targeting are listed in supplemental information (Table S1) were designed using CRISPR design tools from http://crispr.mit.edu:8079 (Zhang laboratory, MIT). The synthesized oligo DNA primers for the guides were inserted into pX330 (Addgene #42230). For Tet-inducible expression of exogenous TopoII, TopoII cDNAs fused with mCherry at their N-terminus were inserted into MluI/SalI site on the hH11 Tet-ON cassette (Natsume et al., 2016) donor plasmid described previously (Hassebroek et al., 2020). cDNA fragments of TopoIIα and TopoIIβ CTD were amplified from full length cDNA and then cloned in the pET3 0a plasmid (EMD Millipore/Novagen). The TopoIIα truncation mutants were generated using PCR, the ChT swapping mutants using fusion PCR and the point mutations using site-directed mutagenesis by a QuikChangeII kit (Agilent) with the primers listed in supplemental information (Table S1). All constructs were verified by DNA sequencing. The parental line construct for OsTIR1 targeting and the TopoIIα-AID construct were created as described previously (Hassebroek et al., 2020).

For preparation of recombinant TopoII-CTD proteins, the proteins were expressed in Rossetta2 (DE3) strain with previously established culture condition (Ryu et al., 2010). The bacterial pellet was harvested the following day and the cells were lysed using lysozyme in Lysis Buffer (450 mM NaCl, 30 mM HEPES (pH 7.7), 0.5 mM TCEP). After incubation at 4°C for 1h, 5% Glycerol, 1% Triton-X 100, 10 units/ml DNase1,10 mM MgCl_2_, 0.1 mM PMSF and 0.5 mM TCEP were added, and the suspension was incubated again for 1h at 4°C. The suspension was then centrifuged at 12000 rpm at 4°C for 30 minutes. His6-tagged protein in the supernatant was captured on Talon Sepharose beads (#635502, Clontech/Takara) and pre-equilibrated against Buffer1 (300 mM NaCl, 20 mM HEPES (pH 7.7), 2 mM MgCl_2_, 2.5 mM Imidazole and 0.5 mM TCEP). After incubation with Talon beads at 4°C for 1-2h, the beads bound with protein were emptied into a column, the column was washed with 5 volumes of Buffer1 containing 2.5 mM ATP, 5 mM MgCl_2_ and 2.5 mM Imidazole and 0.5 mM TCEP and 2 volumes of Buffer2 (50 mM NaCl, 10 mM HEPES (pH 7.7) and 2 mM MgCl_2_) containing 2.5mM Imidazole and 0.5mM TCEP. Elution was then carried out using 15 mM, 7 5 mM, and 450 mM Imidazole in Buffer 2. The eluted fractions were then tested using SDS-PAGE followed by CBB staining. The fractions that contained the protein were pooled and subjected to Hi-trap ion exchange chromatography (GE healthcare) for further purification.

The α-PICH and α-TopoIIα antibodies were generated as previously described (Hassebroek et al., 2020). TopoIIβ-antibody was kindly provided by K. Tsutsui (Tsutsui et al., 2001).

### Cell culture and Transfections

CRISPR/Cas9 targeted insertion was performed as previously described to generate all the cell lines (Hassebroek et al., 2020). In brief, DLD-1 cells were transfected with the guide and donor plasmids using Viafect (#E4981, Promega) reagent. Transfections were set up in 3.5 cm dishes. 2days after, the cells were trypsinized and replated on a 10cm dish at ≈20% confluency and starting from Day 3, they were subjected to a selection process by maintaining them in the presence of a suitable selection reagent (1μg/ml blasticidine [#ant-bl, Invivogen], 0.5μg/ml Puromycin [#ant-pr, Invivogen], 200μg/ml hygromycin B gold [#ant-hg, Invivogen]). After 10-14 days of this process, the colonies were isolated and cultured in 48-well plates. The colonies were subjected to Western Blotting and Genomic PCR analyses to verify the integration of the transgene. For Western Blotting analyses, the cells were pelleted and boiled/vortexed with 1X SDS-PAGE sample buffer. The samples were analyzed using antibodies as described in each figure legend.

Genomic DNA was isolated by cell pelleting and lysis using lysis buffer (100 mM Tris-HCl (pH 8.0), 200 mM NaCl, 5 mM EDTA, 1% SDS, and 0.6 mg/ml proteinase K [#P8107S, NEB]) followed by ethanol precipitation and resuspension with TE buffer containing 50 μg/ml RNase A (#EN0531, ThermoFisher). The obtained genomic DNA samples were subjected to PCR using primers indicated in the supplemental information to ensure integration at the correct locus.

The TopoII AID cell lines were established using the OsTIR1 parental DLD-1 line (Hassebroek et al., 2020). The DNA coding for the AID-3XFLAG tag was integrated into the TopoIIα or TopoIIβ locus using CRISPR/Cas9 editing in the parental cells as indicated in the supplemental information. The candidate clones obtained were screened by genomic PCR to verify accurate transgene integration. Once validated for integration, the ability of Aux to deplete the protein was tested by Western blotting and Immunostaining. The mCherry-TopoII wt/ mutant replacement cell lines were also engineered using CRISPR/Cas9 in the TopoIIα-AID cell line by inserting the gene coding for the rescue candidate at the hH11 locus (Zhu et al., 2014), as previously described (Hassebroek et al., 2020) and as indicated in the supplemental information. The isolated clones were validated for transgene integration using genomic PCR analysis. The expression of the mCherry-fusion protein was confirmed using western blot upon addition of Dox. The CENP-A-miRFP670/mNeon-TopoIIα line is created using OsTIR/AID-PICH line (Hassebroek et al., 2020), then validated as shown in supplemental figure 7 with genomic PCR.

### Preparation of Whole Cell Lysate and Mitotic Chromosome Samples

For preparation of the whole cell lysate for testing most of AID system-based degradation, cells were synchronized with thymidine for 18-24h. Upon release, the cells were replenished with media containing 0.5 mM Aux to allow degradation of endogenous AID-tagged proteins. After ≈ 8.5 hours, the cells were then harvested and boiled/vortexed with 1X SDS-PAGE sample buffer and subjected to western blotting with antibodies as indicated in the figure legends. For preparation of mitotic chromosomes from the TopoIIα-AID and TopoIIβ-AID cell lines, a mitotic shake-off was performed. For this, the cells were plated and at ≈ 70% confluency, they were treated with thymidine. After 18 hours, the cells were released from thymidine for 6 hours following which they were treated with nocodazole (100ng/ml) for 4 hours. The mitotic cells were then released from the plate, washed using 1X McCoy’s devoid of FBS, three times, and resuspended in fresh 1X McCoy’s media containing FBS for releasing them from nocodazole for 30 minutes. Post nocodazole release, the cells were pelleted and incubated on ice for 5 minutes with lysis buffer (250 mM sucrose, 20 mM HEPES (pH 7.7), 100 mM NaCl, 1.5 mM MgCl_2_, 1 mM EDTA, 1 mM EGTA, 0.2% Triton X-100, 1:2000 LPC [leupeptin, pepstatin, chymostatin; 20 mg each/ml in DMSO; Sigma-Aldrich], and 20 mM iodoacetamide [Sigma-Aldrich #I1149]). Chromosomes were isolated from the lysed cells by loading the lysate on to 40% glycerol cushion and centrifuging at 3000 rpm for 5 minutes at 4°C. After washing with 40% glycerol cushion, the isolated chromosomes were boiled and vortexed with 1X SDS-PAGE sample buffer and subjected to western blotting.

The whole cell lysates for verifying induction of the mCherry-fusion protein with doxycycline were prepared by synchronizing cells with thymidine (-Aux/+Aux /+Aux+Dox samples were all consistently prepared). Upon release from thymidine block, the cells were concurrently treated with 0.5 mM Aux and 100 μg/ml Dox to allow degradation of endogenous TopoIIα with the simultaneous expression of the rescue candidate, ≈8.5 hours after which the cells were harvested and subjected to western blotting as specified in the above two scenarios.

The primary antibodies used were α-TopoIIα (in-house), α-TopoIIβ (Tsutsui et al., 2001), α-PICH (in-house), α-FLAG M2 (#F1804, Millipore/Sigma), α-mCherry (#ab167453, Abcam), α-H4 (#61521, Active Motif) and β-tubulin (#T-4026, Sigma). β-tubulin was used as the loading control for whole cell lysate samples and α-H4 was used in the case of Chromosome samples.

### Western Blotting

The samples for SDS-PAGE were loaded and separated on handmade/gradient (Invitrogen/ThermoFisher scientific) gels followed by transfer on to a methanol activated PVDF membrane using an ECL-semidry transfer unit (Amersham Biosciences). Following blocking using casein (for higher molecular weight proteins) or gelatin (for lower molecular weight proteins), the proteins of interest were selectively probed for using primary antibodies as indicated in each figure legend. The secondary antibodies used were IRDye 800CW secondary antibodies and IRDye 680 RD antibodies (LICOR). Signals were visualized using the LI-COR Odyssey Fc machine.

### UFB assays and Immunostaining

DLD-1 cells were grown for no longer than 10 passages in McCoy’s 5A 1X L-Glutamine containing 7.5% Fetal Bovine serum (FBS). Cells were plated on coverslips coated with Poly-D-Lysine (#A38904-01, gibco/Life Technologies Corporation) in a 3.5 cm dish. The coating was done by incubating the flame sterilized coverslips in Poly-D-Lysine solution for >1h. The coat was then removed, and the coverslips were allowed to dry for >3h. Cells were incubated with 2mM Thymidine (added at ≈ 70% confluency) for 18-24h for synchronization. Thymidine was washed off using McCoy’s media lacking FBS three times. After washing off Thymidine, 0.5mM Auxin and 1μg/ml Doxycycline were added to media containing FBS and the cells were incubated in the same to allow release from the Thymidine treatment. ≈8.5 hours post-release, a major fraction of cells “roundup”. At this point, the cells were fixed and stained using suitable antibodies. For Fig.1C and Fig.1D, cells for analyses were obtained by performing a mitotic shake-off, after which the cells were washed and released from the nocodazole arrest for 30 minutes to 1 hour to allow progression from the prometaphase arrest. The cells were then plated on fibronectin coated coverslips (#GG-12-1.5-Fibronectin, NEUVITRO) that allows the mitotic cells to be retained on them. The cells were fixed and stained as follows. Fixation was performed using a 4% solution of paraformaldehyde (pFA) in 1X PBS for 10 minutes. After washing off the pFA, the cells were permeabilized using methanol at −20°C for 10 minutes. The cells were then blocked with a 2.5% solution of hydrolyzed gelatin prepared in 1X PBS containing 0.1% Tween-20 (1X PBS-T) for 1 hour. Following this, the cells were incubated with a cocktail of primary antibodies for 3 hours, washed with 1X PBS-T for 10 minutes, thrice, and then treated with secondary antibodies for 1h before being washed and mounted onto slide glasses with VECTASHIELD Antifade Mounting Medium containing 4’,6-diamidino-2-phenylindole (#H-1200, Vector Laboratory) and sealed using nail polish.

For scoring the UFB phenotype, we focused on anaphase cells at a consistent stage by ensuring that the CENP-C signals between the segregating chromosomes were at a similar distance, as well as after making sure that the DAPI signals were well distinguishable between segregating chromosomes. Cells which had progressed to anaphase-B were excluded from the counting procedure. PICH staining was used as a readout for UFBs as PICH localizes to these structures (Baumann et al., 2007; Ke et al., 2011; Spence et al., 2007). All the UFBs that can be visualized were counted by switching the plane of imaging. The UFB number for each of the cells was recorded. For each cell line represented in this study, the UFB number was obtained from a minimum of 60 cells visualized across three independent experiments with at least 20 cells being counted from each experiment.

The stained cell images were acquired using the Plan Apo 100x/1.4 objective lens on a Nikon TE2000-U equipped PRIME-BSI CMOS camera (Photometrics) with MetaMorph imaging software. Figures were prepared from exported images by adjusting the intensity with Image J software, following the guideline of the Journal.

For quantification of TopoIIα on mitotic chromosomes in live cells, endogenous TopoIIα was depleted, and mCherry-tagged alleles were induced as described above. To ensure cells were at an equivalent stage of mitosis, nocodazole was added 20 minutes before imaging. Then, mCherry in live cells was imaged under normal cell culture conditions, at 37°C and 5% CO_2_, with nocodazole, using a DeltaVision Ultra microscope fitted with an Olympus 60X/1.42, Plan Apo N objective (UIS2, 1-U2B933), C-Y-R Polychroic, and PCO-Edge sCMOS camera (>82% QE), using the following acquisition parameters. Entire mitotic cell volumes were obtained by capturing 24μm thick Z-series with 0.2μm spacing, i.e. 120 slices per cell. 50ms exposure and 10% lamp power resulted in a rapid Z-series capture time of approximately 6 seconds, which limited image distortion that would have resulted from chromosome movements. Z-series were cropped above and below the cell then projected using SoftWoRx software, then mCherry signal was quantified using ImageJ software by averaging the signal intensities across a 50-pixel wide line spanning the chromosomes.

The antibodies used were α-PICH (Hassebroek et al., 2020), rat α-DYKDDDDK (#200474-21, Agilent) for FLAG signals, 5F8 α-RFP (#RMA5F8, Bulldog Bio), α-CENP-C (#PD030, MBL), goat α-rabbit IgG Alexa Fluor 488 (#A11034, Molecular probes/Life Technologies), goat α-rat IgG Alexa Fluor 568 (#A11077, Molecular probes/Life Technologies, Invitrogen), goat α-guinea pig IgG Alexa Fluor 647 (#A21450, Molecular probes/Life Technologies).

### Morphological quantification of TopoII distribution with wndchrm

Images for wndchrm analyses were acquired from live cells expressing fluorescent TopoIIα, as described above. For quantification of morphological similarities/dissimilarities, a supervised machine learning algorithm, wndchrm (weighted neighbor distance using a compound hierarchy of algorithms representing morphology) was 1.52 was used, as previously described (Matsumoto et al., 2016; Ono et al., 2017; Takagi et al., 2018; Tokunaga et al., 2014). For the mCherry-TopoIIα wt and its mutant proteins in Fig. 4 D-E, 30 images for each kind were collected, and randomly subdivided into two subgroups. For supplemental figure S6, 6, 8, and 10 images of mCherry-TopoIIα and mCherry-TopoIIα Y1521A were used. For Fig. 5 G, 30, 40, 50, 60 and 70 images of TopoIIα in the control and GSK-343 treated cells were used. Morphological feature values of the image were automatically assigned by training a machine. For each test, cross-validation tests were automatically repeated 20 times using 70% of images as training and 30% of images as test among the provided data set. The options used for wndchrm analysis were a large feature set of 2919 (-l) and multi-processors (-m). Morphological distances between two classes (class A and class B) were calculated as the Euclidean distances [d=√∑(A–B)2] with the values in a class probability matrix obtained from the cross validations. P values were also provided by two-sided Student’s t-tests for each of the comparisons. Phylogenies were created with the PHYLIP package ver. 3.67 (Felsenstein, 1989; Johnston et al., 2008), using pairwise class similarity values computed by wndchrm.

### Mono-nucleosome Pull Down assays

Recombinant S-tagged TopoII-CTD proteins were loaded on S-protein Agarose beads (#69704, EMD Millipore/ Novagen) by incubating them together overnight. Log phase chromatin was isolated from DLD-1 cells after cell lysis using lysis buffer (as stated in the previous section for chromosome isolation) and centrifugation after loading on with 20% glycerol cushion. The isolated chromatin was salt extracted using high salt buffer (400 mM NaCl, 18 mM β-Glycerophosphate, 20 mM HEPES (pH 7.7), 5 mM EDTA, 5mM EDTA and 5% glycerol) and digested in Micrococcal Nuclease (MNase) buffer (50 mM NaCl, 20 mM HEPES (pH 7.7), 5% glycerol, 5 mM CaCl_2_) with MNase (#M0247S, New England Biolabs). The obtained digest (mononucleosomes) was incubated with the beads coated with the proteins for 1h at 22°C. The beads were then washed with 1X TBS-T containing 250 mM NaCl and the results were analyzed using western blotting. Additionally, a fourth of the beads were treated with half-diluted salt extraction buffer containing 10% SDS and 0.5μl Proteinase-K. The samples were run on a DNA gel and the size of the input and bound digest were verified. Representative input and bound digest are shown in supplemental figure 3.

The primary antibodies used for analyses included α-H3K27me2 (#ab24684, abcam), α-H3K27me3 (#C36B11, Cell Signaling Technology), α-H3K27me3 (#61018, Active Motif), α-H3 (#96C10, Cell Signaling Technology), α-H4 (#61521, Active Motif). For visualizing the S-tagged bait, S-protein HRP conjugate (#69047, EMD Millipore/Novagen) was used followed by chemiluminescence substrate Pico PLUS (#34577, Thermo Scientific Protein Biology).

The western blot signals were visualized using the LI-COR Odyssey Fc machine and the band intensities were analyzed using the Image Studio Lite software. Each of the band intensities for H3K27me3 signal were normalized with respect to the S-HRP (bait signal) for the corresponding sample. The intensity of H3K27me3/bait values was then recorded in Graphpad prism software using which statistical analyses were performed.

### Optimization of GSK-343 addition conditions and assays with GSK-343 treatment

TopoIIα-AID cells engineered with hH11-Tet-ON-mCherry-TopoIIα wt were used for optimization assays. These cells were used without any Aux or Dox treatment. GSK-343(#A3449, APExBIO) was diluted to 6 mM, 4 mM, and 2 mM concentration from the stock in DMSO medium. The protocol employed is the same as the UFB assay protocol with 1:1000 GSK-343 was added from each of the abovementioned dilutions to obtain 2 μM, 4 μM and 6 μm final concentration of the inhibitor. The inhibitor was added with wash off of the cells from Thymidine. Equal amounts of DMSO was added to control cells. Subsequently 8.5h later, the cells were fixed with 4% pFA, permeabilized with methanol and stained for H3K27me3 (#C36B11, Cell Signaling Technology), PICH (inhouse antibody) and CENP-C (#PD030, MBL), goat α-rabbit IgG Alexa Fluor 488 (#A11034, Molecular probes/Life Technologies), goat α-rat IgG Alexa Fluor 568 (#A11077, Molecular probes/Life Technologies), goat α-guinea pig IgG Alexa Fluor 647 (#A21450, Molecular probes/Life Technologies). DNA was visualized using DAPI.

Once the inhibitor concentration was optimized for our experimental conditions, 4 μM final GSK-343 was used in assays with endogenous TopoIIα depletion and exogenously expressing mCherry TopoIIα-wt/mutants. This time, the inhibitor was diluted to 16mM concentration from the stock. With removal of thymidine and the consequent addition of Aux and Dox, 1:4000 GSK-343 (from 16 mM) was added to obtain final concentration of 4 μM. This higher dilution factor was chosen to further reduce the amount of DMSO added to the cells along with the inhibitor (since Aux and Dox are also diluted in DMSO). Equal amounts of DMSO were added to control cells. The cells were then similarly fixed and stained as stated above with antibodies were α-PICH (Hassebroek et al., 2020), 5F8 α-RFP (#RMA5F8,Bulldog Bio), α-CENP-C (#PD030, MBL), goat α-rabbit IgG Alexa Fluor 488 (#A11034, Molecular probes/Life Technologies), goat α-rat IgG Alexa Fluor 568 (#A11077, Molecular probes/Life Technologies), goat α-guinea pig IgG Alexa Fluor 647 (#A21450, Molecular probes/Life Technologies).

## Statistical analyses

The quantifiable data was analyzed for statistical significance using GraphPad Prism software (version 8). Based on the data itself, either the one-way ANOVA or paired two-tailed t-test was employed followed by suitable post-hoc tests where applicable.

## Acknowledgements

We thank K. Tsutsui and M. Miyaji for the anti-TopoIIβ antibody. This work was supported by NIH/NIGMS, GM112793 and GM130858, then in part, by KUCC/CB pilot grant (KAN1000623) and General research funds from University of Kansas (#2144098 and #2144083). It was also supported by JSPS KAKENHI Grant Numbers JP18H05531, JP18K19310, JP20H03520 [to N.S.], and by grants from The Vehicle Racing Commemorative Foundation [to N.S.]. A. Arnaoutov and M. Dasso are supported by NIH/NICHD Intramural projects Z01 HD008954 and ZIA HD001902.

## Author Contributions

SS conducted chromatin pull-down assays, created TopoII mutant replaced cell lines in Fig. 3, performed UFB assay with them, optimized GSK-343 treatment condition and performed UFB assays in Figs 6 and 7, and drafted manuscript. HP created AID-TopoIIα cell line and most of TopoII mutant replaced lines, performed UFB assays in Figs 1, 2 and supplemental figure 4, and acquired images. SK established AID-TopoIIβ line then performed initial UFB assay in Fig. 1 for TopoII-depleted cells. DC co-designed study with YA and performed live cell imaging and analysis in Figs 4 and 5 together with MJ. TF and NS performed wndchrm analysis of the images in Figs 4 and 5. AA and MD provided original gene targeting plasmids for OsTIR and CENP-A. YA designed the study, supervised project, and co-wrote the manuscript with DC.

## Supporting Information

**Table S1.**
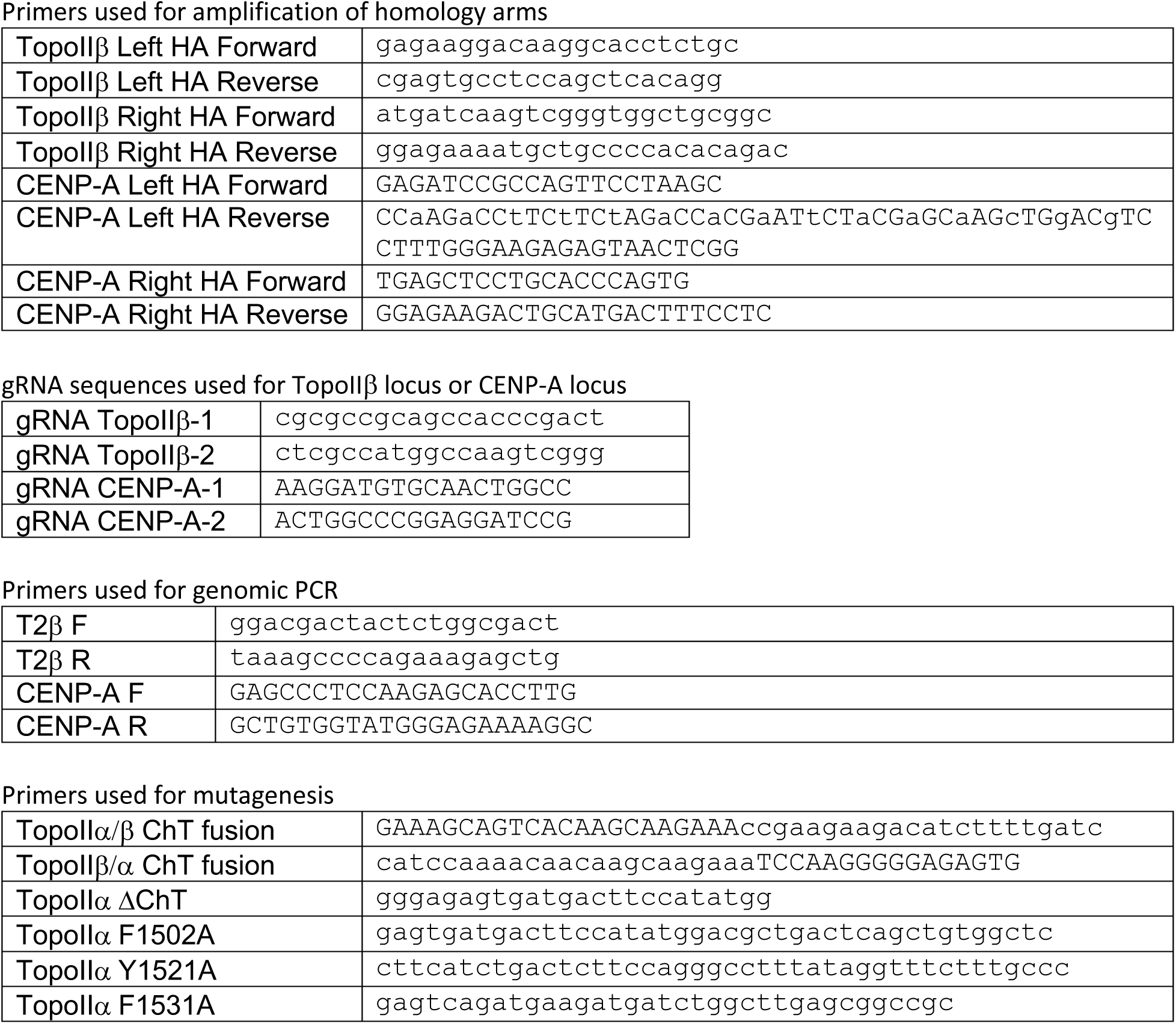

## Abbreviations

AID: Auxin inducible degron
Aux: Auxin
Dox: Doxycycline
CEN: Centromere
ChT: Chromatin Tether
CTD: C-terminal domain
PICH: Polo-like kinase interacting checkpoint helicase
SPR: Strand passage reaction
Tet: Tetracycline
TopoIIα: Topoisomerase IIα
TopoIIβ: Topoisomerase IIβ
UFB: Ultra-fine DNA bridge

---

**Supplemental Figure 1.**
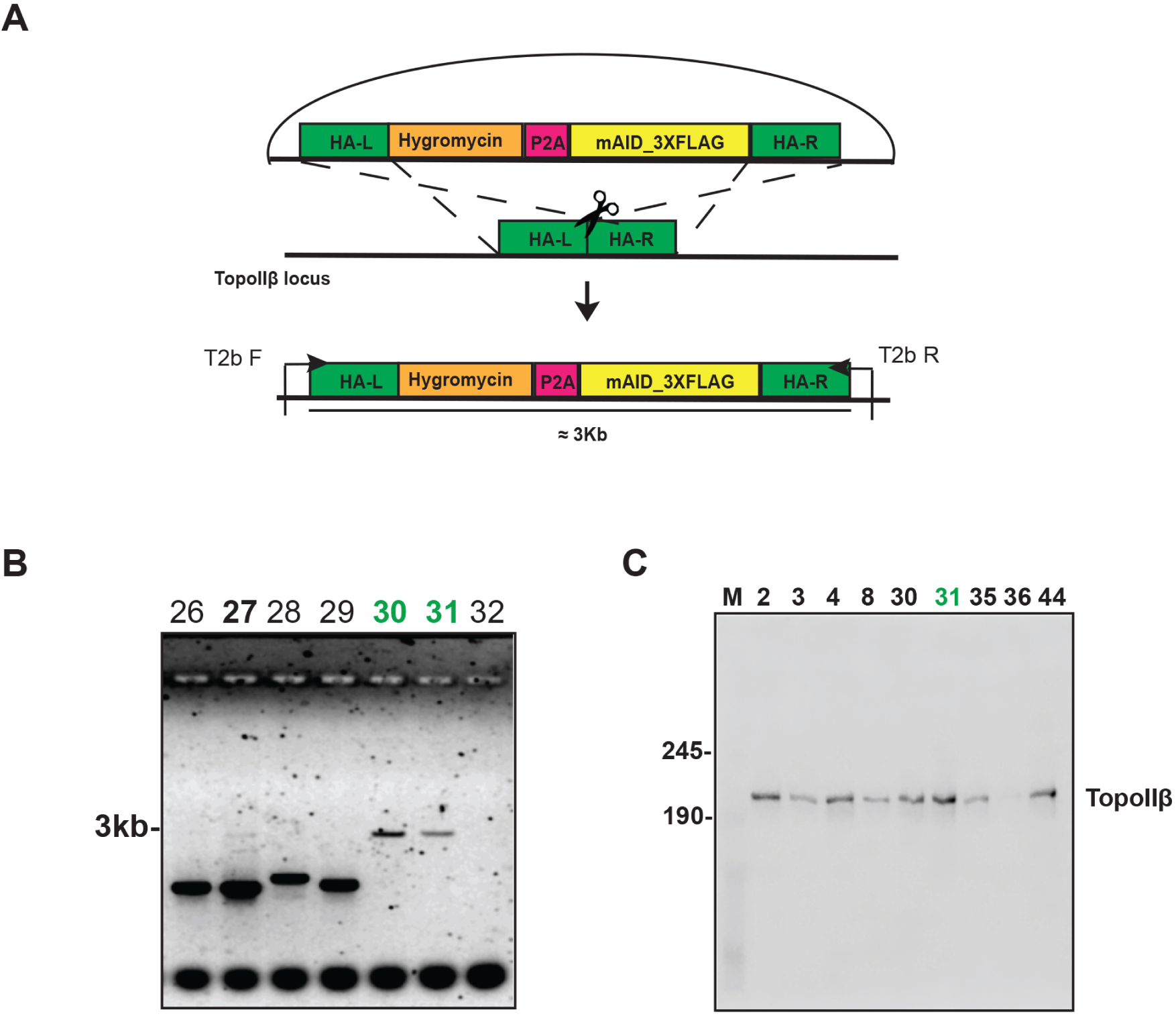
Construction of the TopoIIβ-AID line. **A**-Schematic Representation of donor plasmid for tagging the 5’ end of TopoIIβ with AID. Cells were transfected with the donor plasmid along with two guides. **B**- Representative genomic PCR for the selected Hygromycin resistant clones. Genomic DNA was isolated from a number of clones and PCR was performed using the primers indicated in A. Clones showing the 3kb band only are homozygous AID-integrated (#30, #31) indicated in green **C**- Representative Western Blot of the whole cell lysate obtained from the hygromycin resistant clones. Anti-TopoIIβ antibody was used to detect the tagged TopoIIβ. Among the clones, we chose to work with clone #31

**Supplemental Figure 2:**
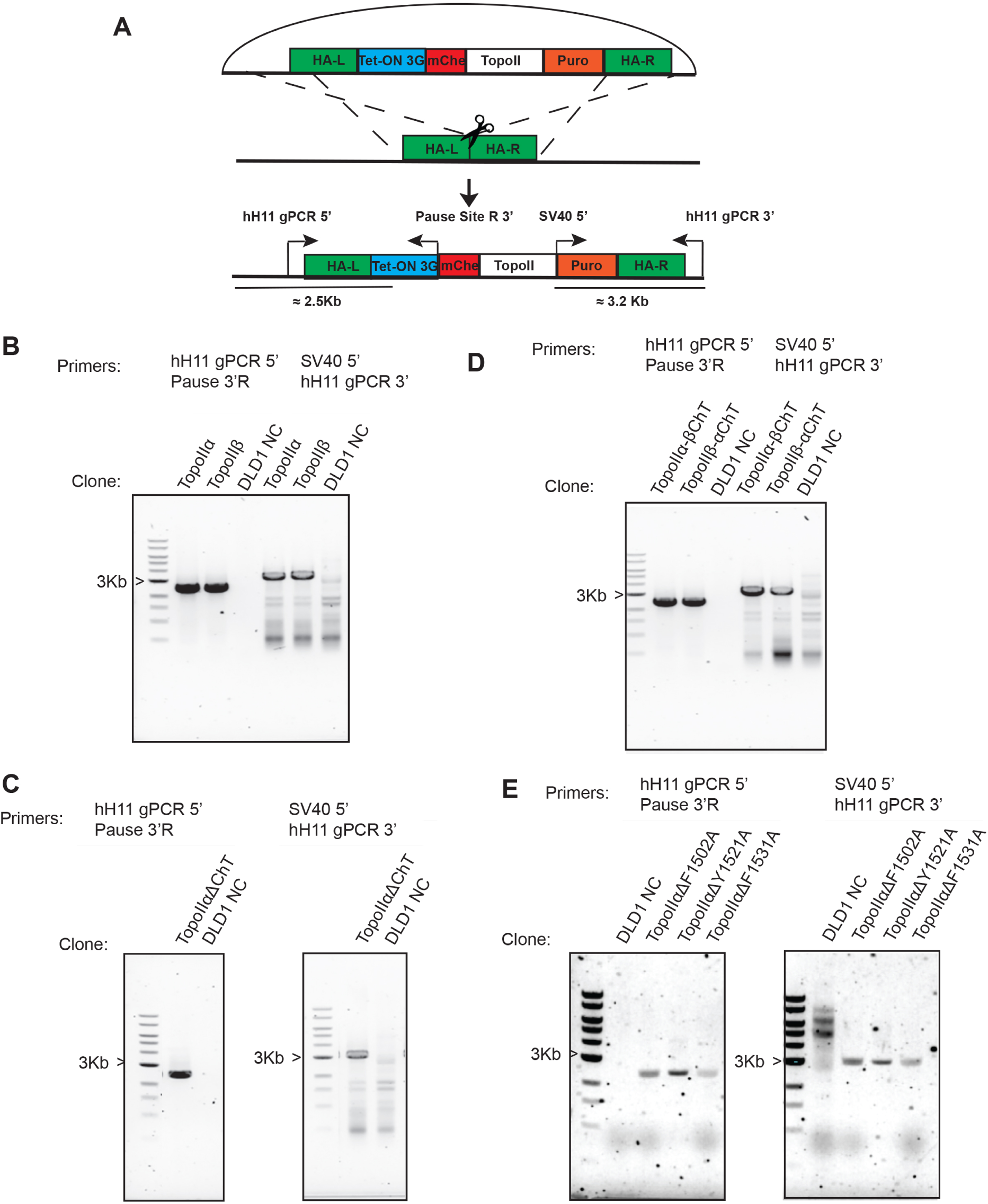
Construction of the hH11 Tet-ON TopoII replacement cell lines. **A**- Schematic Representation of donor plasmid containing the replacement cassette. Cells were transfected with the donor plasmid and two guides for the hH11 locus. Clones were selected for Puromycin resistance. **B/C/D/E**- Representative Genomic PCR for TopoIIα wt and TopoIIβ wt (B), TopoIIα–ΔChT mutant (C), TopoIIα ChT-swapping mutants (D) and aromatic cage mutants (E) replacement. Genomic DNA isolated from the clones were subjected to PCR with primers targeting the 5’ end as well as the 3’ end as indicated in A. Non-transfected DLD-1 cells were used as the control (DLD1-NC).

**Supplemental Figure 3.**
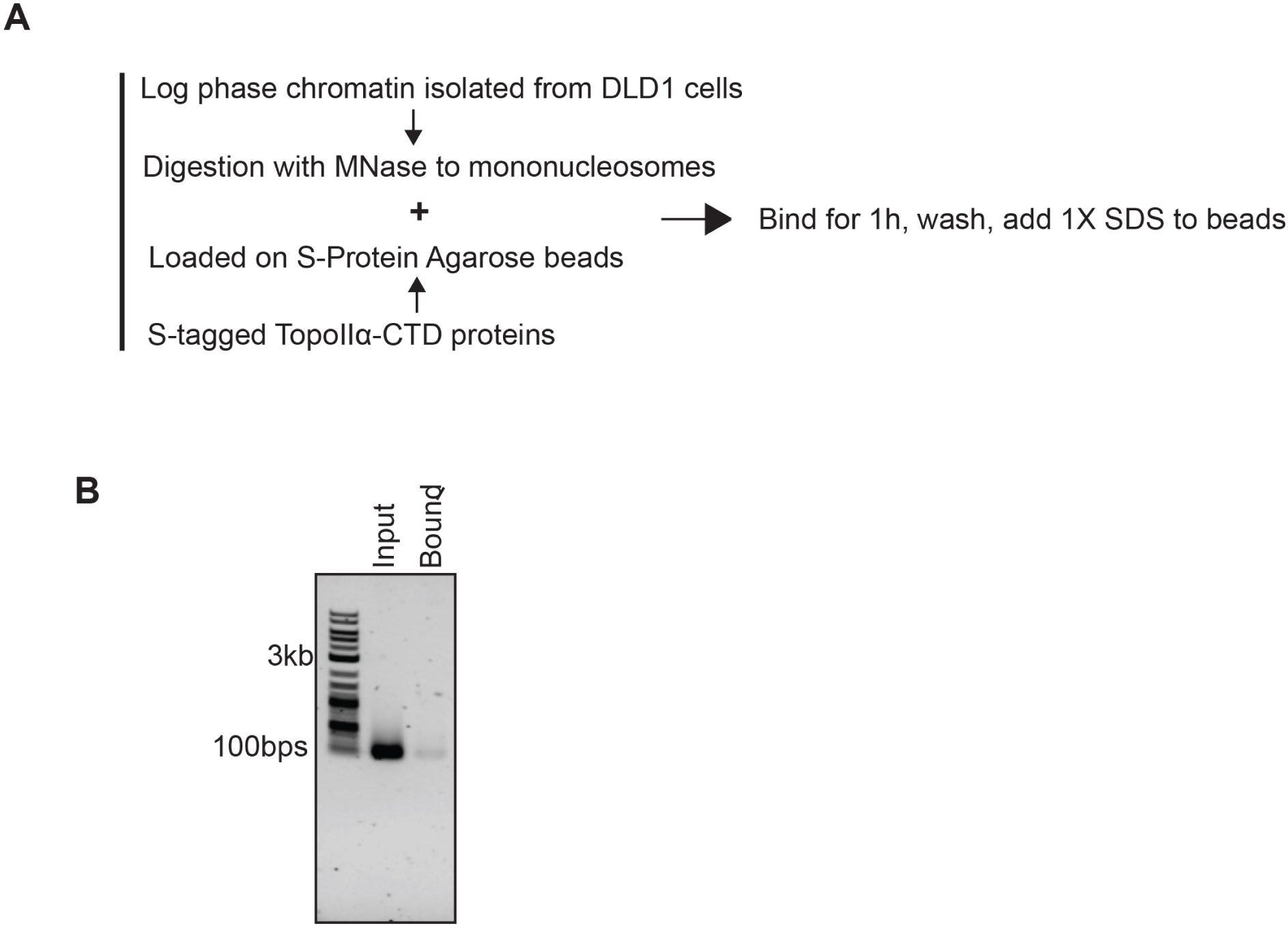
Mononucleosome preparations used in chromatin pull down assays. **A**- Schematic representation of protocol used to perform chromatin pull down assays. **B**- Representative image indicating mononucleosomes after digestion of log phase chromatin isolated from DLD-1 cells after MNase treatment for input (left) and bound to beads (right) samples. The size of the band (≈140bps) indicates that the majority of input and bound chromatins are mononucleosomes.

**Supplemental Figure 4.**
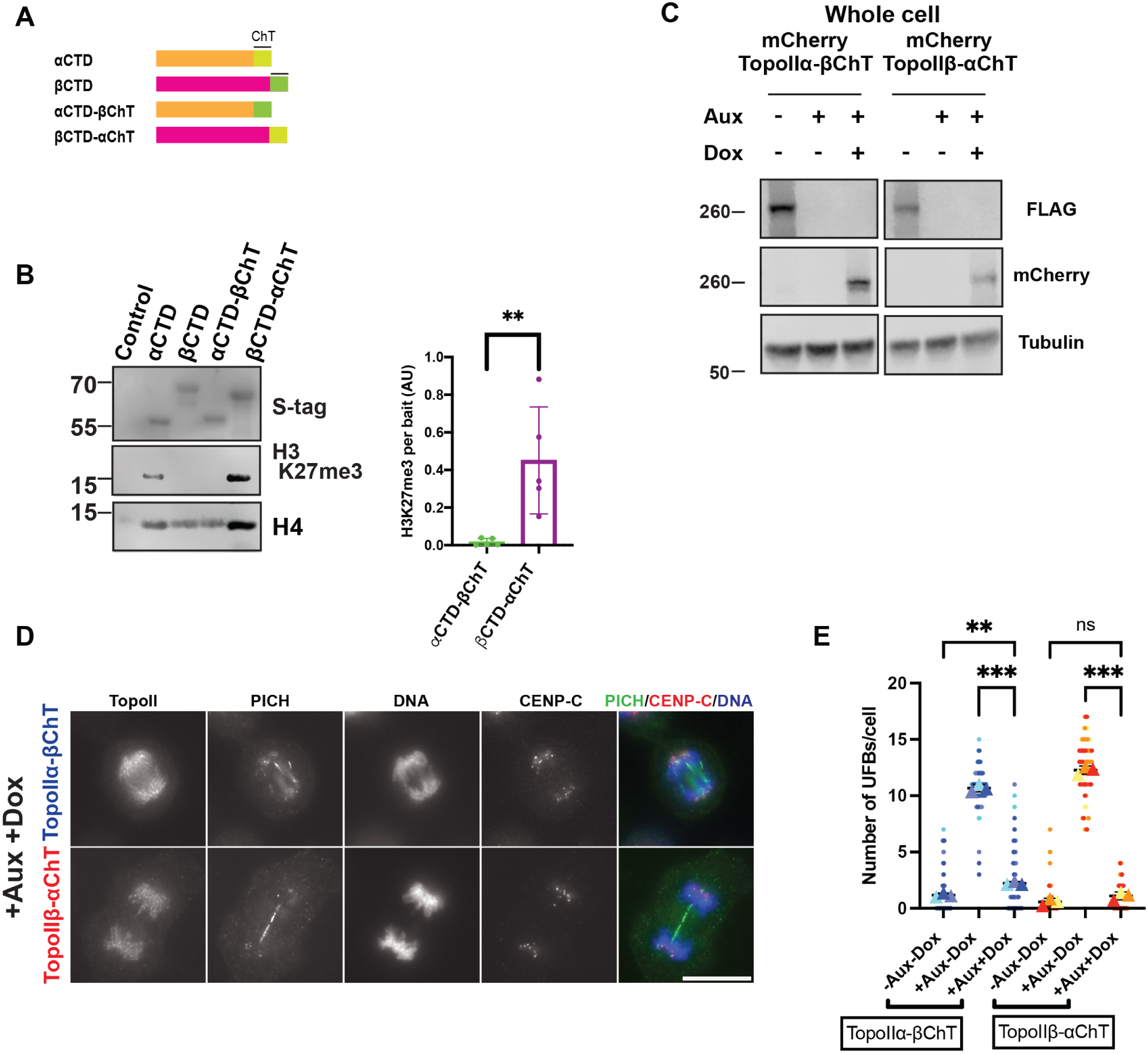
ChT domain swapped mutants highlight the importance of the αChT in H3K27me3 containing chromatin binding and in complete UFB resolution. **A**- Schematic representation of TopoIIα and TopoIIβ ChT domain swapped mutants (top). The terminal 31 amino acids of each of TopoIIα and TopoIIβ were swapped with one another. **B**- Results from chromatin pull down for these mutants indicates that the ChT domain dictates binding to H3K27me3 containing chromatin (bottom left). Quantification of H3K27me3 binding per bait (AU) from N=5 experiments (bottom right). Swapping their ChT domain reverses the ability of TopoIIα-CTD and TopoIIβ-CTD to bind H3K27me3 containing chromatin. P value indicates two-tailed unpaired samples t-test. Error bars indicate SD. *:p<0.033, **:p<0.002, ***:p<0.001. **C**- Western blot representing replacement of the full-length TopoIIα-βChT and TopoIIβ-αChT mutants in endogenous TopoIIα replacement background. **D**- UFB assay for TopoIIα and TopoIIβ ChT domain swapped mutants’ replacement in TopoIIα depleted cells. PICH staining for UFBs indicated that the binding of these mutants to H3K27me3 (Fig 3D) correlates with its ability to resolve UFBs. Bars, 10μm. **E**- Superplots for quantification of the number of UFBs/cell (from >60 cells counted over three independent experiments) for TopoIIα and TopoIIβ ChT domain swapped mutants’ replacement in TopoIIα depleted cells. P-value indicates oneway ANOVA analysis followed by Tukey multicomparison correction. Horizontal bars indicate mean and error bars indicate SD calculated for the means across the three independent experiments. **:p<0.002.

**Supplemental Figure 5.**
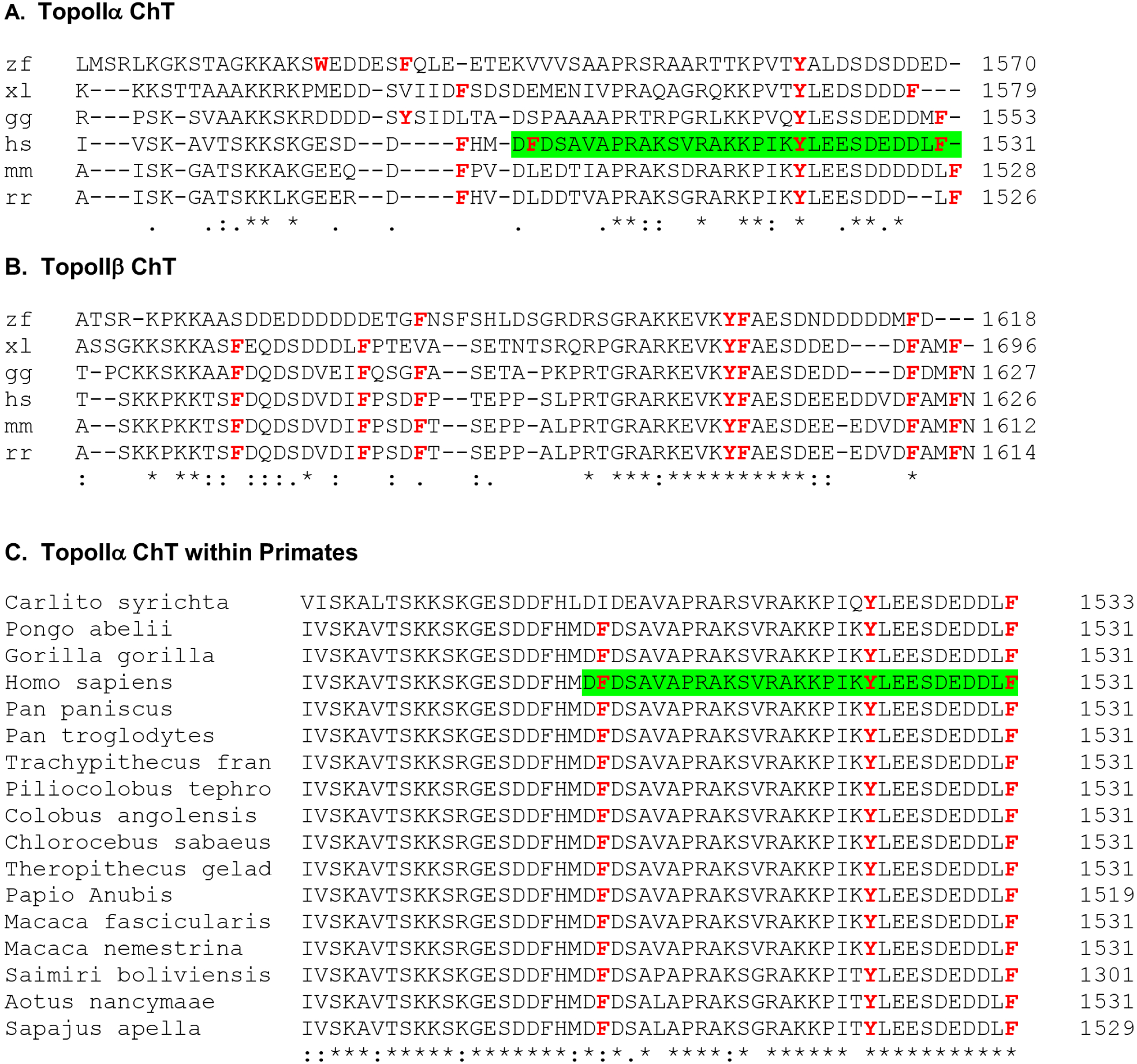
The aromatic aminoacids in the αChT are highly conserved among primates. **A**- Sequence alignment for the TopoIIα-ChT domain among the indicated vertebrates. **B**- Sequence alignment for the region corresponding to the ChT domain in TopoIIβ among the indicated vertebrates. **C**- High levels of conservation of the three aromatic residues-F1502, Y1521 and F1531 in primates indicated by the sequence alignment. For A and B; zf: Zebrafish, *Danio rerio,* xl: African clawed frog, *Xenopus laevis,* gg: Chicken, *Gallus gallus,* hs: human, *Homo sapiens,* mm: Mouse, *Mus musculus,* rr: Rat, *Rattus rattus.* Symbols in the arraignment indicates conservation among compared groups as follow;. weak, : strong, and * identical.

**Supplemental Figure S6.**
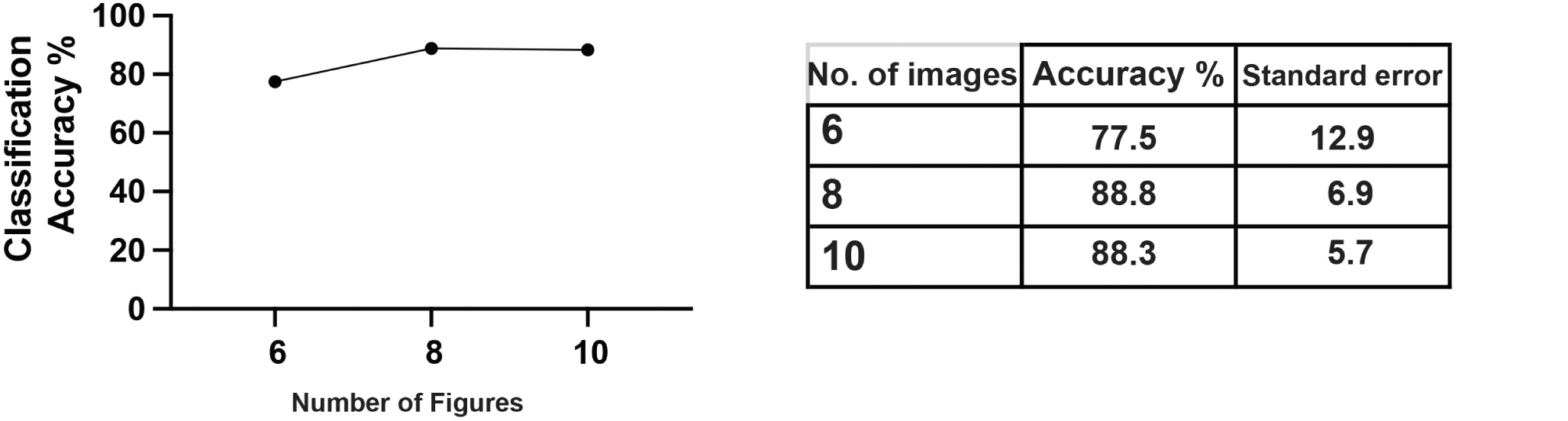
Classification accuracies for discrimination of TopoIIα morphologies. Wndchrm was used to discriminate the localization patterns of TopoIIα -wt and -Y1521A. Increasing numbers of training image sets were used. 10 images were sufficient to achieve accurate classification (88.3%).

**Supplemental Figure 7.**
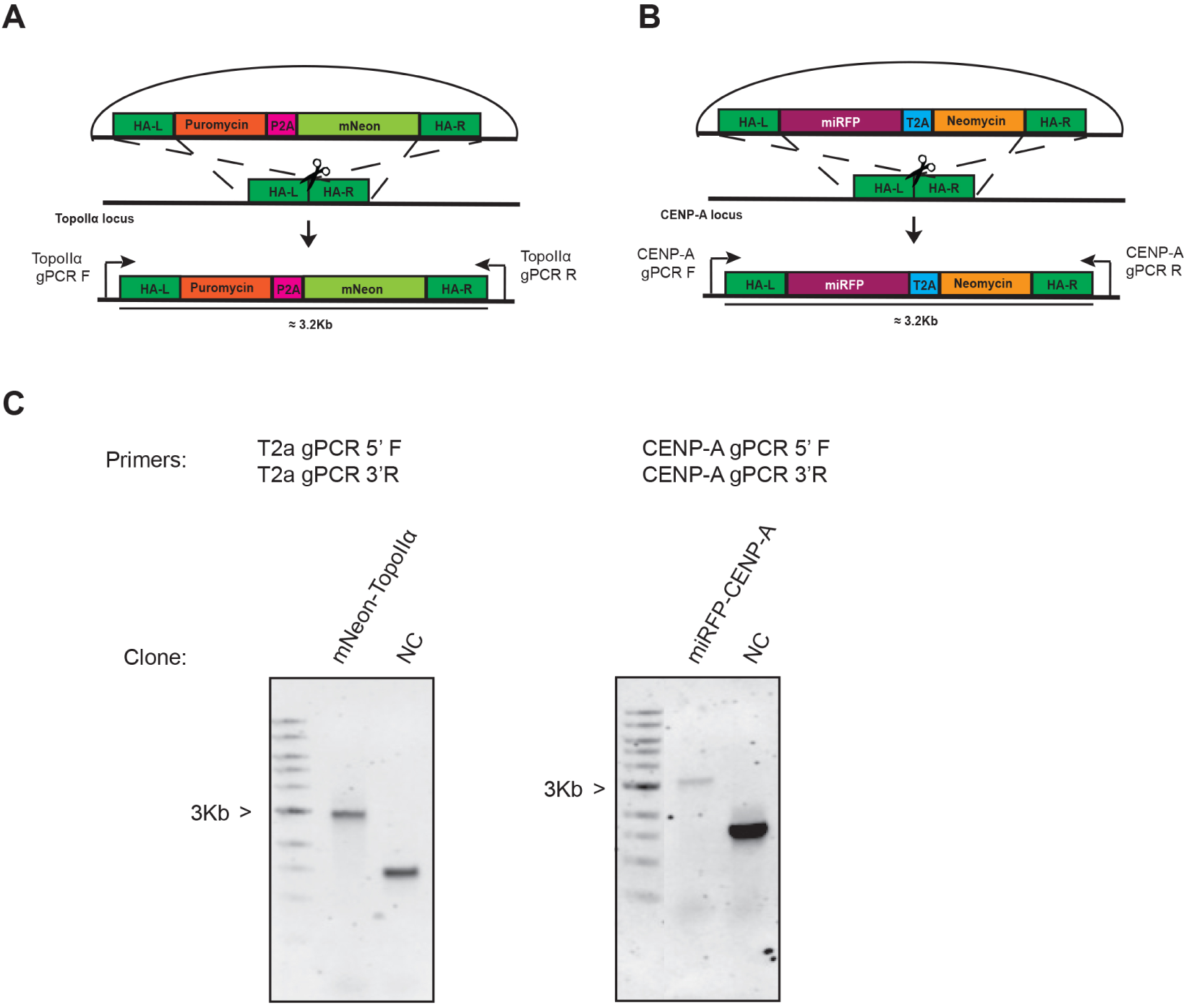
Construction of the mNeon-TopoIIα / CENP-A-miRFP670 line. **A**- Schematic Representation of donor plasmid for tagging the 5’ end of TopoIIα with mNeon. **B**- Schematic Representation of donor plasmid for tagging the 3 ‘ end of CENP-A with miRFP. **C**- Representative Genomic PCR for mNeon-TopoIIα and miRFP-CENP-A. Genomic DNA isolated from the clones were subjected to PCR with primers targeting the 5’ end as well as the 3’ end as indicated in A. Non-transfected DLD-1 cells were used as the control (DLD1-NC).

**Supplemental Figure 8.**
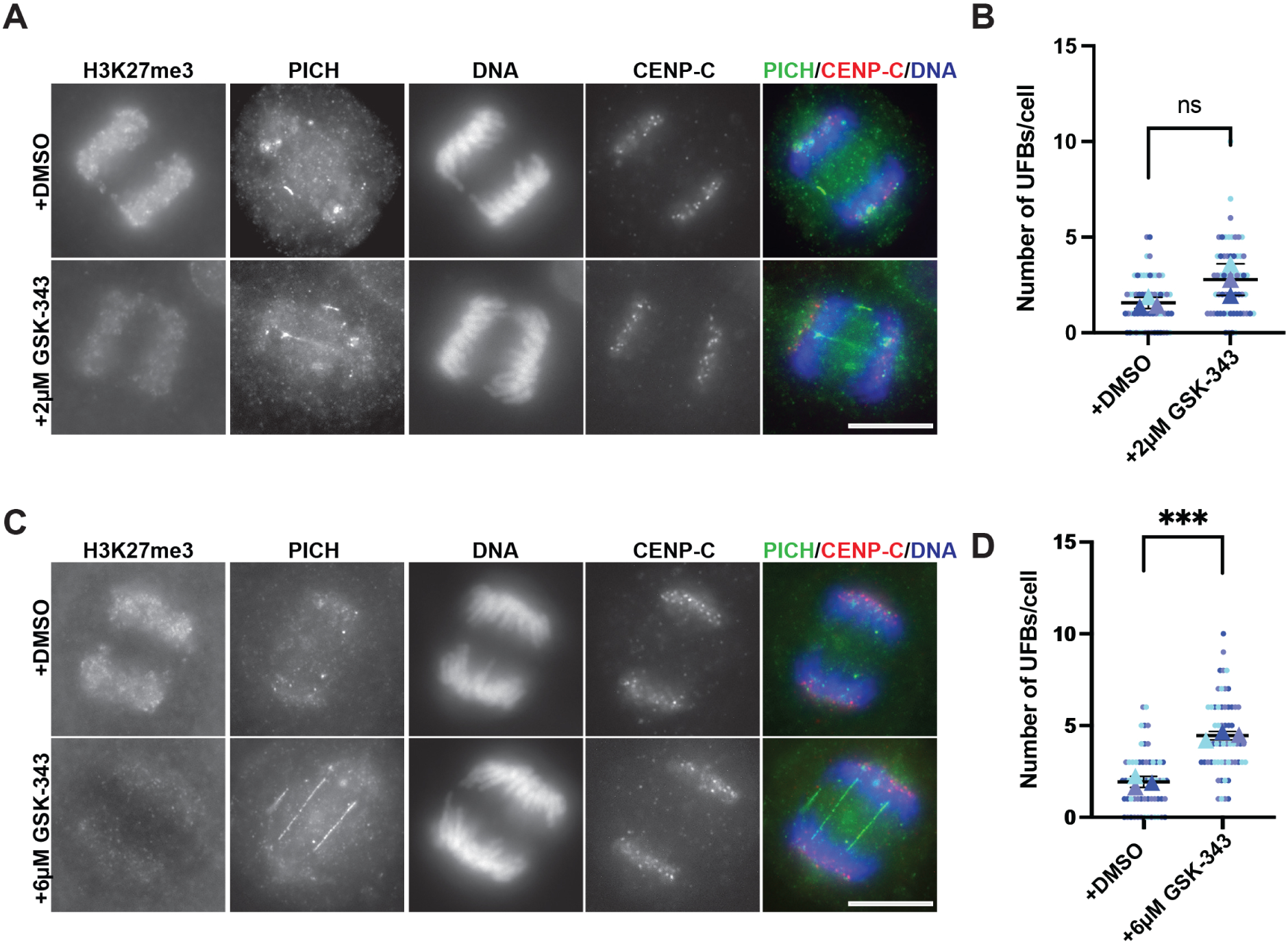
GSK-343 concentration dependency in UFB assays. **A/C**- Representative images from UFB assays with 2 μM (A) and 6 μM (C) GSK-343 and their respective controls (DMSO). Staining with H3K27me3 antibody indicated reduction in signal following GSK-343 treatment. Bars, 10 μm. **B/D**- Superplots showing quantification of the number of UFBs/cell (from >60 cells counted over three independent experiments) for UFB assays with 2μM (B) and 6μM (D) GSK-343 treatment as compared to control cells. P-value indicates one-way ANOVA analysis followed by Tukey multicomparison correction. Horizontal bars indicate mean and error bars indicate SD calculated for the means across the three independent experiments. ns: not statistically significant, ***:p<0.001.

**Supplemental Figure 9.**
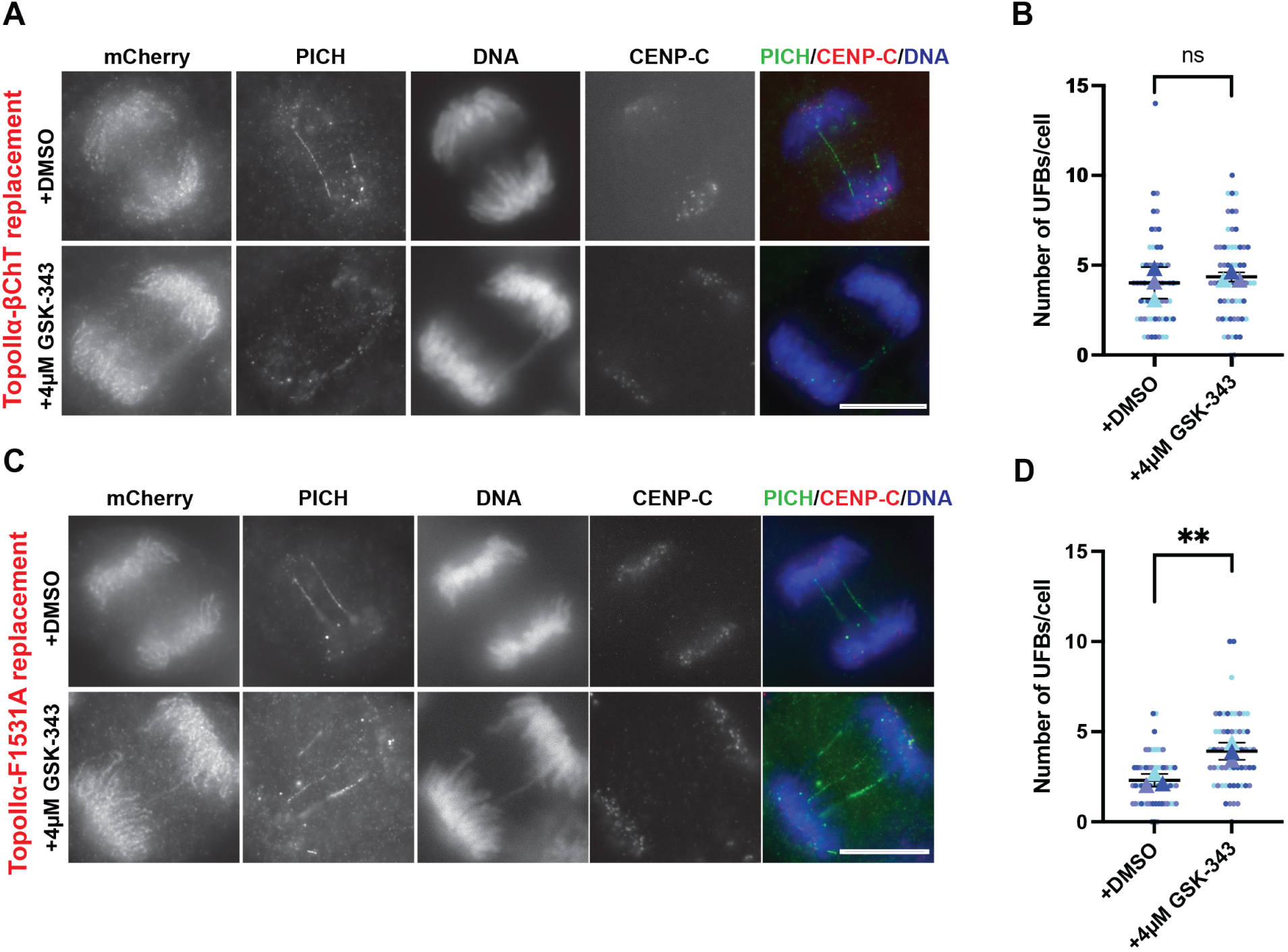
Additional mutants studied for synergistic effect upon H3K27me3 inhibition. **A/C**- Results from UFB assays for mCherry-TopoIIα-βChT (A) and for mCherry-TopoIIα -F1531A (C) replacement in endogenous TopoIIα depleted cells. Upon 4 μM GSK-343 addition, no increase in unresolved UFB number was observed in the former and an increase in unresolved UFB number was observed in the latter as compared to control cells. Bars, 10 μm. **B/D**- Superplots showing quantification of the number of UFBs/cell (from >60 cells counted over three independent experiments) for UFB assays with mCherry-TopoIIα-βChT (B) and mCherry-TopoIIα -F1531A (D) with 4 μM GSK-343 treatment as compared to control cells. P-value indicates one-way ANOVA analysis followed by Tukey multicomparison correction. Horizontal bars indicate mean and error bars indicate SD calculated for the means across the three independent experiments. ns: not statistically significant, **:p<0.002.

## Reference

Adriaens, M.E., P. Prickaerts, M. Chan-Seng-Yue, T. van den Beucken, V.E.H. Dahlmans, L.M. Eijssen, T. Beck, B.G. Wouters, J.W. Voncken, and C.T.A. Evelo. 2016. Quantitative analysis of ChIP-seq data uncovers dynamic and sustained H3K4me3 and H3K27me3 modulation in cancer cells under hypoxia. Epigenetics Chromatin. 9:48.

Antoniou-Kourounioti, M., M.L. Mimmack, A.C.G. Porter, and C.J. Farr. 2019. The Impact of the C-Terminal Region on the Interaction of Topoisomerase II Alpha with Mitotic Chromatin. Int J Mol Sci. 20.

Baumann, C., R. Korner, K. Hofmann, and E.A. Nigg. 2007. PICH, a centromere-associated SNF2 family ATPase, is regulated by Plk1 and required for the spindle checkpoint. Cell. 128:101–114.

Biebricher, A., S. Hirano, J.H. Enzlin, N. Wiechens, W.W. Streicher, D. Huttner, L.H. Wang, E.A. Nigg, T. Owen-Hughes, Y. Liu, E. Peterman, G.J.L. Wuite, and I.D. Hickson. 2013. PICH: a DNA translocase specially adapted for processing anaphase bridge DNA. Mol Cell. 51:691–701.

Chan, K.L., P.S. North, and I.D. Hickson. 2007. BLM is required for faithful chromosome segregation and its localization defines a class of ultrafine anaphase bridges. EMBO J. 26:3397–3409.

Diaz-Martinez, L.A., J.F. Gimenez-Abian, Y. Azuma, V. Guacci, G. Gimenez-Martin, L.M. Lanier, and D.J. Clarke. 2006. PIASgamma is required for faithful chromosome segregation in human cells. PLoS One. 1:e53.

Dickey, J.S., and N. Osheroff. 2005. Impact of the C-terminal domain of topoisomerase IIalpha on the DNA cleavage activity of the human enzyme. Biochemistry. 44:11546–11554.

Earnshaw, W.C., B. Halligan, C.A. Cooke, M.M. Heck, and L.F. Liu. 1985. Topoisomerase II is a structural component of mitotic chromosome scaffolds. J Cell Biol. 100:1706–1715.

Felsenstein, J. 1989. PHYLIP—Phylogeny Inference Package (Version 3.2). Cladistics. 5:164–166.

Gilroy, K.L., and C.A. Austin. 2011. The impact of the C-terminal domain on the interaction of human DNA topoisomerase II alpha and beta with DNA. PLoS One. 6:e14693.

Grue, P., A. Grasser, M. Sehested, P.B. Jensen, A. Uhse, T. Straub, W. Ness, and F. Boege. 1998. Essential mitotic functions of DNA topoisomerase IIalpha are not adopted by topoisomerase IIbeta in human H69 cells. J Biol Chem. 273:33660–33666.

Hassebroek, V.A., H. Park, N. Pandey, B.T. Lerbakken, V. Aksenova, A. Arnaoutov, M. Dasso, and Y. Azuma. 2020. PICH regulates the abundance and localization of SUMOylated proteins on mitotic chromosomes. Mol Biol Cell. 31:2537–2556.

Hengeveld, R.C., H.R. de Boer, P.M. Schoonen, E.G. de Vries, S.M. Lens, and M.A. van Vugt. 2015. Rif1 Is Required for Resolution of Ultrafine DNA Bridges in Anaphase to Ensure Genomic Stability. Dev Cell. 34:466–474.

Jacobs, S.A., and S. Khorasanizadeh. 2002. Structure of HP1 chromodomain bound to a lysine 9-methylated histone H3 tail. Science. 295:2080–2083.

Johnston, J., W.B. Iser, D.K. Chow, I.G. Goldberg, and C.A. Wolkow. 2008. Quantitative image analysis reveals distinct structural transitions during aging in Caenorhabditis elegans tissues. PLoS One. 3:e2821.

Kang, H., M.N. Shokhirev, Z. Xu, S. Chandran, J.R. Dixon, and M.W. Hetzer. 2020. Dynamic regulation of histone modifications and long-range chromosomal interactions during postmitotic transcriptional reactivation. Genes Dev. 34:913–930.

Ke, Y., J.W. Huh, R. Warrington, B. Li, N. Wu, M. Leng, J. Zhang, H.L. Ball, B. Li, and H. Yu. 2011. PICH and BLM limit histone association with anaphase centromeric DNA threads and promote their resolution. EMBO J. 30:3309–3321.

Krupina, K., A. Goginashvili, and D.W. Cleveland. 2021. Causes and consequences of micronuclei. Curr Opin Cell Biol. 70:91–99.

Lane, A.B., J.F. Gimenez-Abian, and D.J. Clarke. 2013. A novel chromatin tether domain controls topoisomerase IIalpha dynamics and mitotic chromosome formation. J Cell Biol. 203:471–486.

Linka, R.M., A.C. Porter, A. Volkov, C. Mielke, F. Boege, and M.O. Christensen. 2007. C-terminal regions of topoisomerase IIalpha and IIbeta determine isoform-specific functioning of the enzymes in vivo. Nucleic Acids Res. 35:3810–3822.

Martins, N.M., J.H. Bergmann, N. Shono, H. Kimura, V. Larionov, H. Masumoto, and W.C. Earnshaw. 2016. Epigenetic engineering shows that a human centromere resists silencing mediated by H3K27me3/K9me3. Mol Biol Cell. 27:177–196.

Martins, N.M.C., F. Cisneros-Soberanis, E. Pesenti, N.Y. Kochanova, W.H. Shang, T. Hori, T. Nagase, H. Kimura, V. Larionov, H. Masumoto, T. Fukagawa, and W.C. Earnshaw. 2020. H3K9me3 maintenance on a human artificial chromosome is required for segregation but not centromere epigenetic memory. J Cell Sci. 133.

Matsumoto, A., C. Sakamoto, H. Matsumori, J. Katahira, Y. Yasuda, K. Yoshidome, M. Tsujimoto, I.G. Goldberg, N. Matsuura, M. Nakao, N. Saitoh, and M. Hieda. 2016. Loss of the integral nuclear envelope protein SUN1 induces alteration of nucleoli. Nucleus. 7:68–83.

Min, J., Y. Zhang, and R.M. Xu. 2003. Structural basis for specific binding of Polycomb chromodomain to histone H3 methylated at Lys 27. Genes Dev. 17:1823–1828.

Mohammad, F., S. Weissmann, B. Leblanc, D.P. Pandey, J.W. Hojfeldt, I. Comet, C. Zheng, J.V. Johansen, N. Rapin, B.T. Porse, A. Tvardovskiy, O.N. Jensen, N.G. Olaciregui, C. Lavarino, M. Sunol, C. de Torres, J. Mora, A.M. Carcaboso, and K. Helin. 2017. EZH2 is a potential therapeutic target for H3K27M-mutant pediatric gliomas. Nat Med. 23:483–492.

Natsume, T., T. Kiyomitsu, Y. Saga, and M.T. Kanemaki. 2016. Rapid Protein Depletion in Human Cells by Auxin-Inducible Degron Tagging with Short Homology Donors. Cell Rep. 15:210–218.

Nielsen, C.F., D. Huttner, A.H. Bizard, S. Hirano, T.N. Li, T. Palmai-Pallag, V.A. Bjerregaard, Y. Liu, E.A. Nigg, L.H. Wang, and I.D. Hickson. 2015. PICH promotes sister chromatid disjunction and co-operates with topoisomerase II in mitosis. Nat Commun. 6:8962.

Nielsen, C.F., T. Zhang, M. Barisic, P. Kalitsis, and D.F. Hudson. 2020. Topoisomerase IIalpha is essential for maintenance of mitotic chromosome structure. Proc Natl Acad Sci U S A. 117:12131–12142.

Nielsen, P.R., D. Nietlispach, H.R. Mott, J. Callaghan, A. Bannister, T. Kouzarides, A.G. Murzin, N.V. Murzina, and E.D. Laue. 2002. Structure of the HP1 chromodomain bound to histone H3 methylated at lysine 9. Nature. 416:103–107.

Nitiss, J.L. 2009. Targeting DNA topoisomerase II in cancer chemotherapy. Nature reviews. Cancer. 9:338–350.

Ono, T., C. Sakamoto, M. Nakao, N. Saitoh, and T. Hirano. 2017. Condensin II plays an essential role in reversible assembly of mitotic chromosomes in situ. Mol Biol Cell. 28:2875–2886.

Ryu, H., G. Al-Ani, K. Deckert, D. Kirkpatrick, S.P. Gygi, M. Dasso, and Y. Azuma. 2010. PIASy mediates SUMO-2/3 conjugation of poly(ADP-ribose) polymerase 1 (PARP1) on mitotic chromosomes. J Biol Chem. 285:14415–14423.

Sakaguchi, A., and A. Kikuchi. 2004. Functional compatibility between isoform alpha and beta of type II DNA topoisomerase. J Cell Sci. 117:1047–1054.

Shamir, L., N. Orlov, D.M. Eckley, T. Macura, J. Johnston, and I.G. Goldberg. 2008. Wndchrm - an open source utility for biological image analysis. Source Code Biol Med. 3:13.

Spence, J.M., H.H. Phua, W. Mills, A.J. Carpenter, A.C. Porter, and C.J. Farr. 2007. Depletion of topoisomerase IIalpha leads to shortening of the metaphase interkinetochore distance and abnormal persistence of PICH-coated anaphase threads. J Cell Sci. 120:3952–3964.

Takagi, M., T. Ono, T. Natsume, C. Sakamoto, M. Nakao, N. Saitoh, M.T. Kanemaki, T. Hirano, and N. Imamoto. 2018. Ki-67 and condensins support the integrity of mitotic chromosomes through distinct mechanisms. J CellSci. 131.

Tokunaga, K., N. Saitoh, I.G. Goldberg, C. Sakamoto, Y. Yasuda, Y. Yoshida, S. Yamanaka, and M. Nakao. 2014. Computational image analysis of colony and nuclear morphology to evaluate human induced pluripotent stem cells. Sci Rep. 4:6996.

Tsutsui, K., K. Tsutsui, K. Sano, A. Kikuchi, and A. Tokunaga. 2001. Involvement of DNA topoisomerase IIbeta in neuronal differentiation. J Biol Chem. 276:5769–5778.

Vanden Broeck, A., C. Lotz, R. Drillien, L. Haas, C. Bedez, and V. Lamour. 2021. Structural basis for allosteric regulation of Human Topoisomerase IIalpha. Nat Commun. 12:2962.

Verma, S.K., X. Tian, L.V. LaFrance, C. Duquenne, D.P. Suarez, K.A. Newlander, S.P. Romeril, J.L. Burgess, S.W. Grant, J.A. Brackley, A.P. Graves, D.A. Scherzer, A. Shu, C. Thompson, H.M. Ott, G.S. Aller, C.A. Machutta, E. Diaz, Y. Jiang, N.W. Johnson, S.D. Knight, R.G. Kruger, M.T. McCabe, D. Dhanak, P.J. Tummino, C.L. Creasy, and W.H. Miller. 2012. Identification of Potent, Selective, Cell-Active Inhibitors of the Histone Lysine Methyltransferase EZH2. ACS Med Chem Lett. 3:1091–1096.

Zhu, F., M. Gamboa, A.P. Farruggio, S. Hippenmeyer, B. Tasic, B. Schule, Y. Chen-Tsai, and M.P. Calos. 2014. DICE, an efficient system for iterative genomic editing in human pluripotent stem cells. Nucleic Acids Res. 42: e34.

